# Transcriptomics data integration for context-specific modeling of Atlantic salmon metabolism: functional evaluation of methods based on metabolic tasks

**DOI:** 10.1101/2022.09.23.509266

**Authors:** Håvard Molversmyr, Ove Øyås, Filip Rotnes, Jon Olav Vik

## Abstract

**Motivation:** Constraint-based models (CBMs) are used to study the metabolic networks of organisms ranging from microbes to multicellular eukaryotes. Published CBMs are usually generic rather than context-specific, meaning that they do not capture metabolic differences between cell types, tissues, environments, or other conditions. However, only a subset of reactions in a model are likely to be active in any given context, and several methods have therefore been developed to extract context-specific models from generic CBMs through integration of omics data.

**Results:** We tested the ability of six model extraction methods (MEMs) to create functionally accurate context-specific models of Atlantic salmon using a generic CBM (SALARECON) and liver transcriptomics data from contexts differing in water salinity (life stage) and dietary lipids. Reaction contents and metabolic task feasibility predictions of context-specific CBMs were mainly determined by the MEM that was used, but life stage explained significant variance in both contents and predictions for some MEMs. Three MEMs clearly outperformed the others in terms of their ability to capture context-specific metabolic activities inferred directly from the data, and one of these (GIMME) was much faster than the others. Context-specific versions of SALARECON consistently outperformed the generic version, showing that context-specific modeling captures more realistic representations of Atlantic salmon metabolism.

## 1 Introduction

Within a cell, a multitude of biochemical reactions convert available nutrients into the energy and building blocks required to maintain vital processes and to grow. Metabolism is this vast network of reactions, and the emergence of complete genome sequences and high-throughput experimental technologies has enabled scientists to study it on the system level (Nielsen, 2017). Specifically, by identifying and functionally annotating genes in genomes and connecting them to reactions through gene-protein-reaction (GPR) relationships, the complete genome-scale metabolic network of an organism can be outlined in a constraint-based model (CBM) (Heirendt *et al*., 2019; Fang *et al*., 2020). The scope and availability of CBMs has increased greatly over the past few decades, largely thanks to databases of metabolic reactions and models (Norsigian *et al*., 2020; Moretti *et al*., 2021) and methods for automated reconstruction of microbial metabolic networks from genomes (Mendoza *et al*., 2019). Today, CBMs are readily available for many organisms ranging from microbes to multicellular eukaryotes (Gu *et al*., 2019), and a central goal is to integrate these models with omics data to improve predictions (Ramon *et al*., 2018; Noor *et al*., 2019).

The mathematics of CBMs are based on the stoichiometric matrix, in which columns represent reactions, rows represent metabolites, and each entry is the stoichiometric coefficient of a metabolite in a reaction. The other key ingredient is a flux vector that represents the rates of all reactions in the network. Because metabolism is much faster than other biological processes with which it interacts, e.g., transcription and translation, metabolite concentrations are assumed to be constant and a steady-state constraint is imposed on the system (Shamir *et al*., 2016). Along with bounds and any other linear constraints on fluxes, this defines a space of infinitely many solutions, each of which is a feasible combination of fluxes at steady state. This space demarcates an organism’s achievable metabolic states, and thus phenotypes, in a particular environment (O’Brien *et al*., 2015). Optimal states can be found by optimization methods such as flux balance analysis (FBA) (Orth *et al*., 2010), or the whole solution space can be explored through random sampling (Schellenberger and Palsson, 2009) or pathway analysis (Zanghellini *et al*., 2013).

A typical CBM contains intracellular biochemical reactions, transport reactions that move metabolites between cellular compartments, and boundary reactions that allow metabolite exchange with the environment. Furthermore, it is common to add a biomass reaction that allows modeling of growth by accounting for the required molecular building blocks and energy. Maximal growth rate, i.e., maximal flux for the biomass reaction, is often used as the primary objective function for FBA, but any linear combination of reaction rates in the model can be minimized or maximized (Orth *et al*., 2010). Notably, parsimonious FBA (pFBA) simply maximizes growth rate before minimizing total flux (Lewis *et al*., 2010a) but outperforms methods that integrate CBMs with transcriptomics data for flux prediction (Machado and Herrgård, 2014).

Despite the poor performance of transcriptomics-based methods for flux prediction, metabolic activities do differ between contexts such as cells or tissues, and these activities ultimately do depend on upstream processes such as gene expression (Uhlén *et al*., 2015). Generic CBMs, which aim to include all metabolites and reactions found in any cell of an organism, are therefore likely to be superfluous when analyzing specific conditions of interest. Many methods address this by using transcriptomics or other omics data to extract context-specific models rather than to predict or infer fluxes. Context-specific CBMs aim to represent metabolism under a particular set of conditions and have been shown to be more accurate than a generic human CBM (Opdam *et al*., 2017).

The utility of context-specific modeling has been demonstrated through several applications. For instance, tissue-specific CBMs extracted from a genome-scale human CBM have been used to study host-pathogen interactions (Bordbar *et al*., 2010) and brain metabolism (Lewis *et al*., 2010b) as well as for drug target discovery in cancer (Frezza *et al*., 2011). Moreover, context-specific plant CBMs have been used to study fluxes in mesophyll and bundle sheath cells of C_4_ grasses during photosynthesis (Dal’Molin *et al*., 2010), the metabolic behavior of organs for production, storage, and consumption of sugars during the generative phase of barley (Grafahrend-Belau *et al*., 2013), and stress responses to drought in thale cress (Siriwach *et al*., 2020). However, context-specific modeling has not been applied to non-mammalian animals such as fish.

Many different model extraction methods (MEMs) have been developed, employing diverse strategies to create context-specific models by reducing a generic CBM (Robaina Estévez and Nikoloski, 2014). Commonly used MEMs include MBA (Jerby *et al*., 2010), mCADRE (Wang *et al*., 2012), FASTCORE (Vlassis *et al*., 2014), iMAT (Shlomi *et al*., 2008; Zur *et al*., 2010), INIT (Agren *et al*., 2012), and GIMME (Becker and Palsson, 2008), which can be categorized into the MBA-like, iMAT-like and GIMME-like families (Robaina Estévez and Nikoloski, 2014) (**Table 1**). These six MEMs, including their settings, have been thoroughly evaluated for human modeling (Opdam *et al*., 2017; Richelle *et al*., 2019a,b), but the results do not necessarily translate to other animals.

**Table 1.** Description of the six MEMs, grouped by family, with required inputs as implemented in the COBRA Toolbox (Heirendt et al., 2019) and settings used in this study based on recommendations from human studies (Opdam et al., 2017; Richelle et al., 2019a,b).

| Family | Method | Description | Required inputs | Recommended settings |
| --- | --- | --- | --- | --- |
| MBA-like | MBA | Two sets of core reactions are defined with high and medium probability of being active in a given context. MBA then builds a context-specific model containing all high-confidence reactions, as many medium-confidence reactions as possible, and a minimal set of other reactions from the generic CBM to achieve model consistency. | Two sets of core reactions: one with high probability and one with medium probability of being active in the given context. | All reactions associated with a gene score above the 75<sup>th</sup> percentile of the distribution of all gene scores were added to the high confidence set, while all remaining reactions with a score above $5 \ln(2)$ were added to the medium confidence reaction set. The biomass reaction was included in the high-confidence set. |
|  | mCADRE | Using a defined set of core reactions, non-core reactions are pruned based on expression level, connectivity to the core, and a confidence score. Reactions that are not needed to support the core or defined functionalities are removed. If a core reaction is supported by a certain number of unexpressed reactions, it is removed. | Two sets of reaction scores: ubiquity scores quantifying how often each gene is expressed across samples and confidence scores based on evidence from literature. | As the expression distribution of genes is used in the calculation for the gene scores, the gene scores were used as the ubiquity scores. The biomass reaction was given a confidence score of 3. All other reactions were given a score of 1 if they were associated with at least one gene or 0 otherwise. |
| | FAST-CORE | A set of core reactions that should be active in a certain context of interest is defined, and the algorithm finds a minimal set of reactions that support the core. | Single set of core reactions. | All reactions with a gene score greater than $5 \ln(2)$ were added to the core set along with the biomass reaction. |
| iMAT-like | iMAT | Maximizes the number of matches between a reaction's minimal flux and the group it belongs to, i.e., highly or lowly expressed. Thus, it finds a trade-off between including highly expressed reactions and removing lowly expressed reactions. | Two thresholds defining unexpressed and expressed genes along with the gene expression values. | Gene scores were used as expression values and the upper and lower thresholds were both set to $5 \ln(2)$. The biomass reaction was assigned a score of $10 \ln(2)$. |
| | INIT | Finds an optimal trade-off between including and removing reactions based on weights. | A weight for each reaction, which is positive or negative for highly or lowly expressed reactions, respectively. | Reactions with gene score below $5 \ln(2)$ were given a weight of $-8$. Gene score divided by $5 \ln(2)$ was used as weight for other reactions. The maximum weight was used for the biomass reaction. |
| GIMME-like | GIMME | Removes reactions associated with an expression level below a user-defined threshold. Subsequently, reactions are reinserted to achieve a required metabolic function. | A gene expression dataset and an objective function. | Gene scores were used as expression values and the upper and lower thresholds were both set to $5 \ln(2)$. The biomass reaction was assigned a score of $10 \ln(2)$. |

In this study, we used the six MEMs with recommended settings to build context-specific models from SALARECON, a generic CBM of Atlantic salmon metabolism (Zakhartsev *et al*., 2022), using liver transcriptomics data contrasting life stages and feeds (Gillard *et al*., 2018). We evaluated the context-specific CBMs in terms of their contents and predictions, most importantly their ability to perform metabolic tasks (Richelle *et al*., 2019b, 2021), as well as required computation time. Our results show that context-specific model contents and predictions depend heavily on the choice of MEM, but three MEMs produced models that captured significant differences between life stages and one of these was much faster than the others. Context-specific CBMs consistently outperformed SALARECON, supporting context-specific modeling as an approach to explain omics data and improve predictions.

## 2 Methods

### Transcriptomics data and template model

Transcriptomics data from Atlantic salmon liver were downloaded from the GenoSysFat project page on FAIRDOMHub (Wolstencroft *et al*., 2017) (https://fairdomhub.org/projects/34) along with corresponding sample metadata. A detailed description of the feeding trial can be found in the original publication (Gillard *et al*., 2018). Briefly, Atlantic salmon fry were reared in freshwater tanks and continuously fed one of two diets: a feed based on vegetable oil (VO; a combination of linseed oil and palm oil) or a feed based on marine oil (FO; North Atlantic fish oil). About half of the fish were sampled in freshwater and saltwater, respectively. At each life stage, i.e.. either before or after smoltification upon transfer from freshwater to saltwater, about half of the fish were subjected to a feed switch. The most recent version of SALARECON (Zakhartsev *et al*., 2022) was downloaded from its GitLab repository (https://gitlab.com/digisal/salarecon) in February 2021. This version contains the same reactions and metabolites as the published version of SALARECON, but it has 29 fewer genes.

### Extracting context-specific models

Six different MEMs were used to extract context-specific models from the transcriptomics data and SALARECON: MBA (Jerby *et al*., 2010), mCADRE (Wang *et al*., 2012), FASTCORE (Vlassis *et al*., 2014), iMAT (Shlomi *et al*., 2008; Zur *et al*., 2010), INIT (Agren *et al*., 2012), and GIMME (Becker and Palsson, 2008). We used implementations of these MEMs from the COBRA Toolbox (Heirendt *et al*., 2019) to extract context-specific models with the function *createTissueSpecificModel*. The implementation of mCADRE did not perform as expected: if removing a reaction led to an infeasible solution, the procedure stopped with an error. This issue was resolved locally and later merged into the COBRA Toolbox (commit 6c1ba69). The parameters needed to execute the different MEMs were set equal to recommended values (Opdam *et al*., 2017; Richelle *et al*., 2019a,b) (**Table 1**). As the biomass reaction is not directly associated with any genes, steps were taken to ensure its inclusion in all extracted models. Specifically, its lower flux bound was set to a sufficiently small but otherwise arbitrary value of 1 h^*−*1^ for all MEMs. This growth rate was not intended to be realistic and only relative growth rates were used in the analyses. The biomass reaction was also added to the core reaction set of FASTCORE and MBA, assigned a gene score greater than the threshold for GIMME and iMAT, and assigned a specific weight for INIT (**Table 1**).

### Context-specific model contents and predictions

For each context-specific model, we counted the number of genes, reactions, and metabolites, computed maximal growth rate with minimal flux using pFBA (Lewis *et al*., 2010a), and tested the ability to perform metabolic tasks (Richelle *et al*., 2019b, 2021) adapted from mammals to Atlantic salmon (Zakhartsev *et al*., 2022). This was done in Python using COBRApy (Ebrahim *et al*., 2013).

### Gene scores and reaction activity levels

The raw gene expression data was first reduced to only contain genes that were also present in the model. Subsequently, a gene expression threshold was set to determine gene activity in the samples, and any gene with an activity score above this threshold was defined as active. Each gene was given an individual threshold equal to the 90^th^ percentile of its expression value across all samples in the dataset, as this has been documented to yield better models than lower thresholds (Opdam *et al*., 2017). The 25^th^ percentile of the overall gene expression value distribution (i.e., all genes in all samples) was set as the threshold for any gene with a threshold lower than this percentile. A gene score was then computed for each gene (Richelle *et al*., 2019b, 2021):

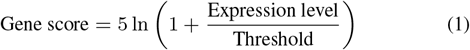

A reaction activity level (RAL) was also computed for each reaction in SALARECON by taking the maximum score of all genes associated with that reaction in the model (Richelle *et al*., 2019b, 2021).

### Metabolic task scores

We obtained a curated and standardized list of 210 metabolic tasks (MTs) covering seven metabolic systems (amino acid, carbohydrate, energy, lipid, nucleotide, and vitamin and cofactor, and glycan metabolism) from the original publication (Richelle *et al*., 2019b). As previously described (Zakhartsev *et al*., 2022), we adapted tasks from mammals to salmon by moving metabolites from compartments not found in SALARECON to the cytoplasm and changing the expected outcomes of amino acid synthesis tasks to match known auxotrophies in salmon. Reactions and associated genes in SALARECON responsible for executing each metabolic task were determined using pFBA (Lewis *et al*., 2010a), and MT scores were calculated as the mean RAL of reactions involved in each task (Richelle *et al*., 2019b, 2021):

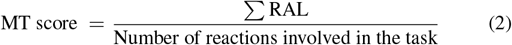

The MT scores were made binary by using the recommended threshold of 5 ln(2) to determine whether or not a metabolic task was performed in a particular sample (Richelle *et al*., 2021). Binary MT scores inferred directly from the data were compared to binary MT scores predicted by context-specific models (task feasibility). For each sample and MEM, we computed the normalized Hamming distance between the MT scores inferred from the data and the MT scores predicted by the model.

### Principal component analysis

Principal component analysis (PCA) was performed to assess the impacts of MEM, feed type and life stage on model contents (reactions) and predictions (metabolic task feasibility). We built two binary matrices in which the rows were models (samples) and the columns were either reactions or tasks. Each column was standardized to have zero mean and unit variance before performing PCA. For each PC, the percentage of variance in PC scores explained by the factors MEM, life stage, and feed was determined by computing the Pearson correlation (*r*) between PC scores and all possible orders of levels for each factor. The maximal *r* across levels was squared to find the explained variance (*R*^2^) for each factor (Opdam *et al*., 2017). We performed PCA once for the whole data set, once for each MEM, and once for each life stage within each MEM.

## 3 Results

### Context-specific model contents and predictions

The transcriptomics data (Gillard *et al*., 2018) covered 81,597 transcripts, 1,109 (14%) of which could be mapped to genes in SALARECON (Zakhartsev *et al*., 2022), across 208 Atlantic salmon liver samples differing in water salinity (life stage) and dietary lipids (feed). There were 112 samples from the freshwater life stage (pre-smolt) and 96 samples from the saltwater life stage (post-smolt), and the fish had been fed diets containing either fish oil (FO) or vegetable oil (VO) with 48 fish switched between feeds at each life stage (FO-VO and VO-FO).

For each sample and each of the six MEMs, we extracted one context-specific CBM using SALARECON. In total, we extracted 1,248 context-specific models, but five mCADRE models were discarded because they were non-functional, leaving 1,243 models. The context-specific CBMs varied significantly in their contents as well as in their predictions (**Fig. 1**). Specifically, models differed between MEMs in terms of gene, reaction, and metabolite counts as well as predicted growth rates, minimal total flux from pFBA (Lewis *et al*., 2010a), and feasible metabolic tasks (Richelle *et al*., 2019b, 2021). Gene, reaction, metabolite, and feasible task counts were highly correlated both between and within MEMs, and larger models tended to predict higher growth rates with less pFBA flux than smaller models (**Fig. S1**). We found no evidence that contents or predictions were significantly affected by life stage (**Fig. S2**) or feed (**Fig. S3**).

**Fig. 1.**
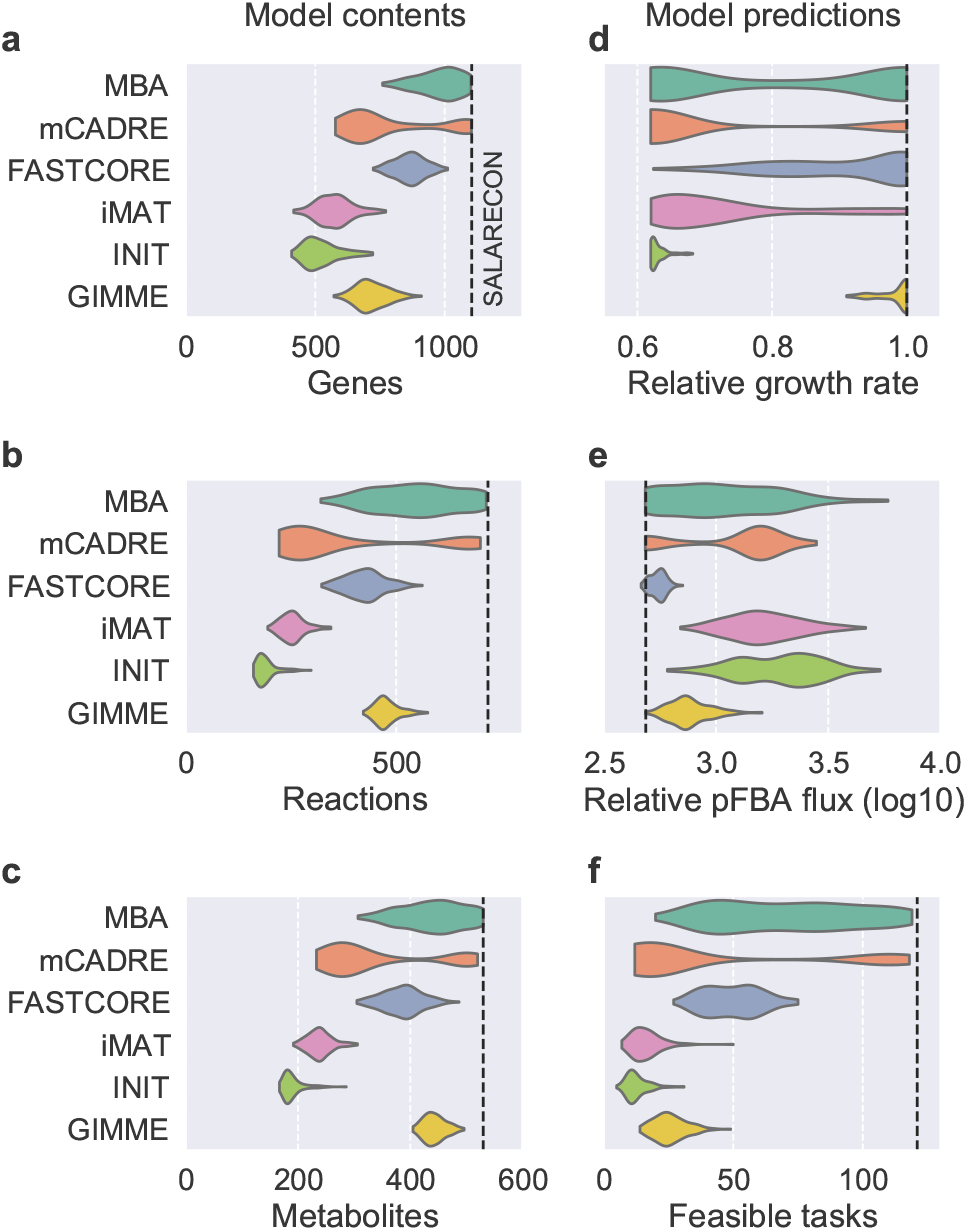
Distribution of context-specific model contents (a–c) and predictions (d–f) by MEM. (a) Gene counts, (b) reaction counts, (c) metabolite counts, (d) predicted maximal growth rate relative to SALARECON, (e) sum of absolute fluxes from pFBA relative to growth rate, and (f) feasible metabolic task counts. Kernel density estimates are scaled to the same width with cutoffs at the extreme data points. Dashed lines indicate predictions from SALARECON.

MBA tended not to reduce the generic model as much as other MEMs, keeping most of the 1,108 genes, 718 reactions, and 530 metabolites from SALARECON but with comparatively wide distibutions for both contents and predictions. Perhaps the most distinguishable MEM was mCADRE, which had bimodal distributions for all contents and predictions: most models were substantially reduced, but some models preserved most of the contents and predictions of SALARECON. Compared to the two other MBA-like MEMs, the contents of FASTCORE models tended to lie between the two mCADRE modes and toward the lower end of the MBA distribution, but FASTCORE predictions were generally more similar to SALARECON. A notable exception is that FASTCORE models performed about half as many metabolic tasks as SALARECON, similar to many MBA and mCADRE models. The two iMAT-like MEMs were very similar across contents and predictions, but iMAT models had a much wider distribution of predicted growth rates than INIT models, all of which had relatively low growth rates. GIMME models were generally most similar to FASTCORE models in their contents, with narrow distributions comparable to the iMAT-like family. Growth rates and minimal flux predictions were close to SALARECON for all GIMME models, but they performed few tasks compared to SALARECON or the MBA-like family.

Looking more closely at the number of context-specific models capable of performing each metabolic task, we found clear differences between tasks, MEMs, and metabolic systems (**Fig. 2**). MBA models tended to perform most of the 121 tasks performed by SALARECON across all systems, reflecting the tendency for MBA to produce models with more reactions than other MEMs. Both MBA and the other MBA-like methods – mCADRE and FASTCORE – had comparatively wide model count distributions, notably spanning the whole range for metabolism of amino acids. The remaining MEMs – iMAT, INIT, and GIMME – were all very similar in terms of the number of models performing tasks. In general, very few models built by these MEMs performed tasks related to metabolism of amino acids, nucleotides, and vitamins, whereas tasks in carbohydrate, energy, and lipid metabolism had a wider range of model counts. The individual tasks that were most frequently performed across all MEMs covered cellular respiration, the thioredoxin system, nucleotide salvage, degradation of ethanol and sugars, as well as synthesis of many amino acids, S-adenosyl methionine (SAM), UDP-glucose, fructose-6-phosphate, glycerol-3-phosphate, and malonyl-CoA (**Fig. S4**). For all MEMs except MBA, there was perfect agreement among models on the feasibility or infeasibility of some tasks. Specifically, all models agreed on two tasks for mCADRE, eight tasks for FASTCORE, 32 tasks for iMAT, 62 tasks for INIT, and 27 tasks for GIMME. Again, we observed no significant effects of life stage (**Fig. S5**) or feed (**Fig. S6**).

**Fig. 2.**
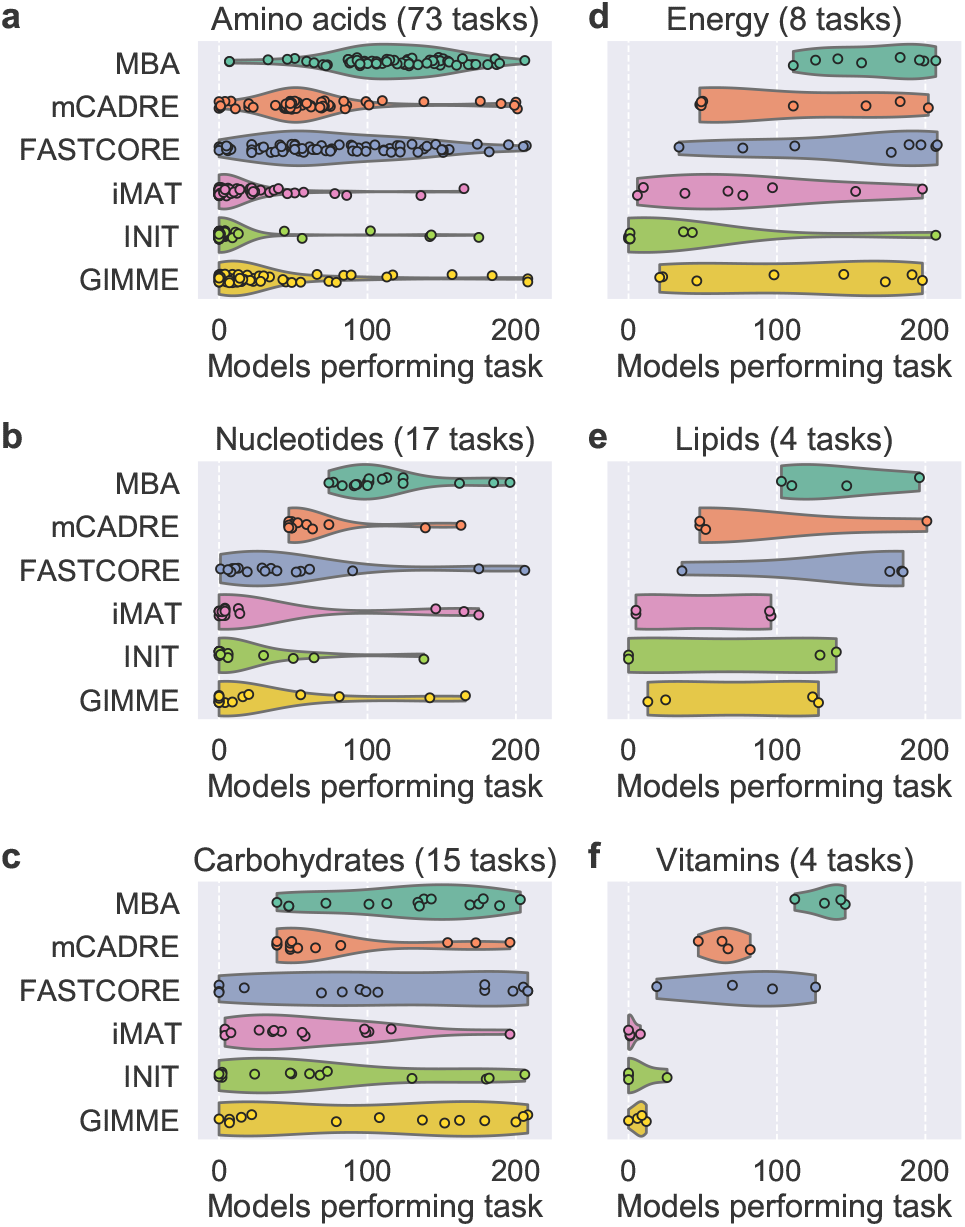
Number of context-specific models in which metabolic tasks are feasible by MEM. Markers represent tasks and are divided into six metabolic systems: (a) amino acid, (b) nucleotide, (c) carbohydrate, (d) energy, (e) lipid, and (f) vitamin metabolism. Kernel density estimates are scaled to the same width with cutoffs at the extreme data points.

### PCA of reaction presence and task feasibility

To disentangle the contributions of MEM, life stage, and feed to the contents and predictions of the context-specific CBMs, we applied principal component analysis (PCA) to binary matrices indicating reaction presence (**Fig. 3**) and metabolic task feasibility (**Fig. 4**). The first two principal components (PCs) explained 38% of the total variance for reactions and 43% of the total variance for tasks, and the scores of models were fairly well-separated by MEM in the first two PCs, both for reactions and for tasks. For both reactions and tasks, the first PC primarily explained variability within mCADRE and MBA models, while the second PC explained more variability between MEMs as well as within FASTCORE, iMAT, INIT, and GIMME models. MEM explained most of the variance of the first five PCs for reactions and the first four PCs for tasks, and these PCs explained almost exactly 50% of the total variance. Each remaining PC explained negligible variance with small contributions from MEM, life stage, and feed (**Figs. S7–S11**).

**Fig. 3.**
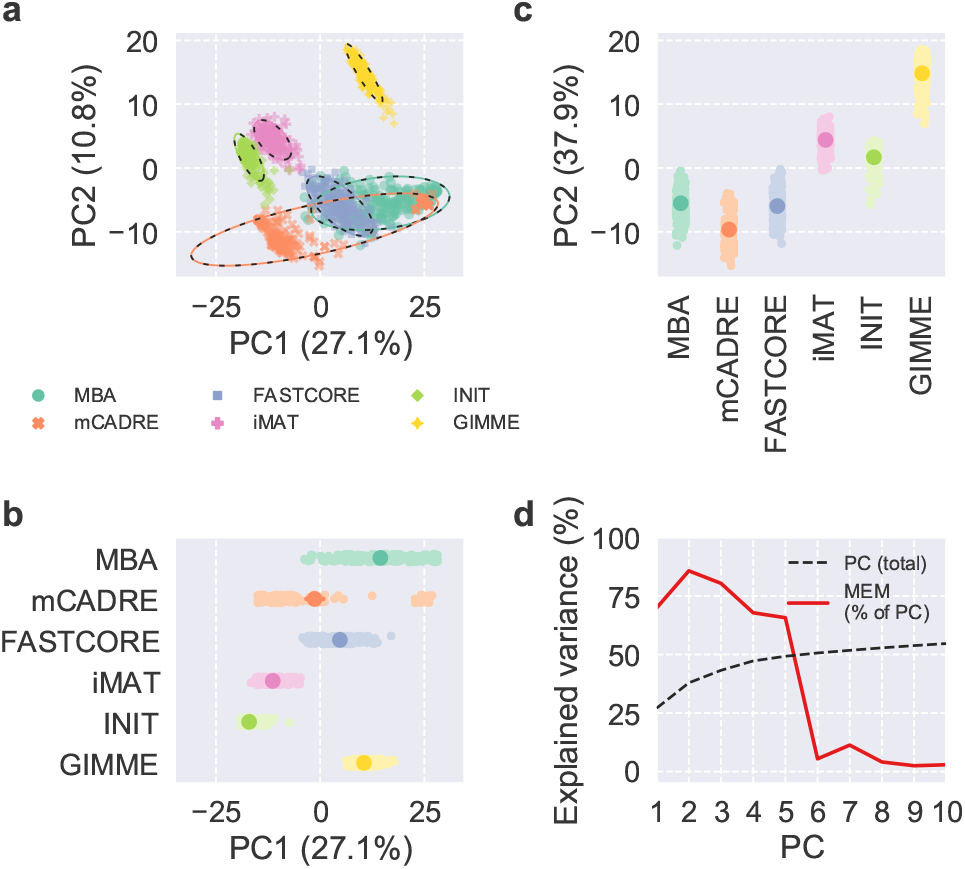
Scores and explained variance from PCA of reaction presence. (a–c) Scores of the first two PCs, colored by MEM, with 95% confidence ellipses and intervals. (d) Cumulative total variance explained by the first ten PCs and variance of PC scores explained by MEM.

**Fig. 4.**
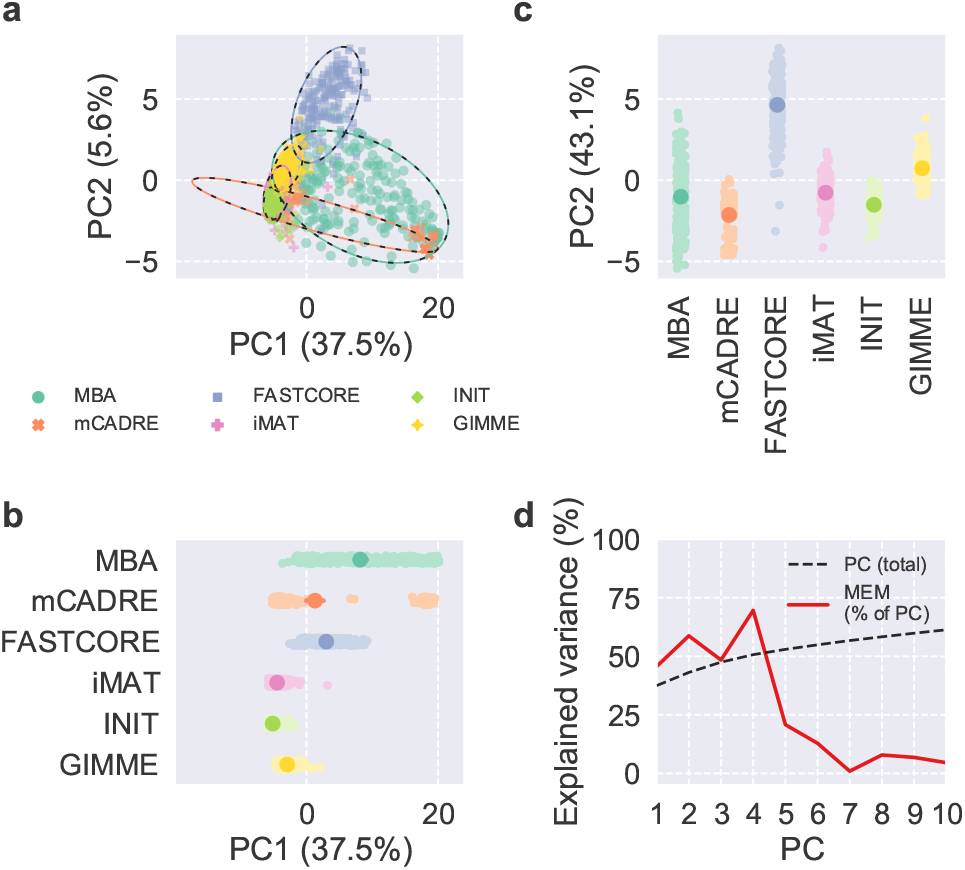
Scores and explained variance from PCA of task feasibility. (a–c) Scores of the first two PCs, colored by MEM, with 95% confidence ellipses and intervals. (d) Cumulative total variance explained by the first ten PCs and variance of PC scores explained by MEM.

Hierarchical clustering of the reaction and task matrices recapitulated the results from PCA with clustering mainly by MEM, but it also seemed to reveal further clustering by life stage within at least some MEMs (**Figs. S12** and **S13**). We therefore applied PCA to reaction presence and task feasibility within each MEM to further interrogate differences between life stages and feeds (**Fig. 5** and **Figs. S14–S37**). For both reactions and tasks, the first PC explained more than 10% variance for MBA, mCADRE, and iMAT, and less than 10% for FASTCORE, INIT, and GIMME. The first PC was dominated outliers for mCADRE and INIT, but INIT’s second PC for tasks also explained more than 10% variance. For three MEMs, life stage explained more than 20% variance for one PC from PCA of reactions: 40% of the first for GIMME, 36% of the second for FASTCORE, and 29% of the third for MBA. For tasks, life stage explained 21% of the second PC for FASTCORE as well as 25% of the second and 22% of the third for GIMME. Feed explained very little variance across all PCs, also within each life stage (**Fig. S38**).

**Fig. 5.**
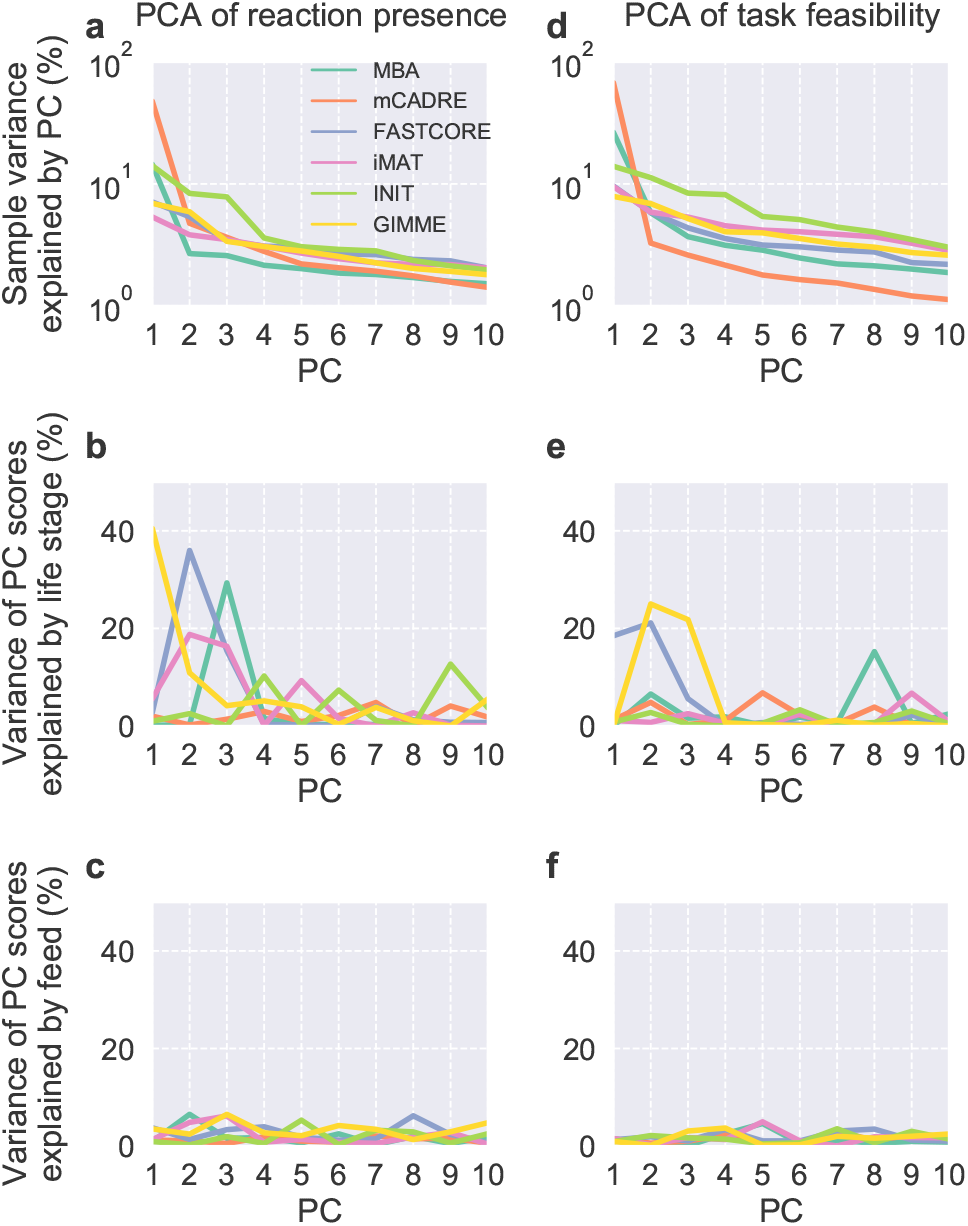
Results from PCA of reaction presence (a–c) and task feasibility (d–f) performed separately for each MEM. Sample variance explained by each PC and variance of PC scores explained by life stage and feed are shown for the first ten PCs.

### Functional accuracy and computational efficiency

To evaluate the functional accuracy of the MEMs, we compared metabolic task feasibility predicted by each extracted context-specific CBM to binary task scores inferred directly from the transcriptomics data that was used for extraction (Richelle *et al*., 2019b, 2021) (**Figs. 6** and **S39**). Specifically, we computed normalized Hamming distances between task feasibility predicted by the models and task scores inferred from the data. We found that iMAT, INIT, and GIMME outperformed the other MEMs in terms of functional accuracy, meaning that they tended to produce models with predicted task feasibility similar to the inferred task scores. In general, MBA models were the least functionally accurate, with a wide distribution covering larger distances than observed for most iMAT, INIT, and GIMME models. Most models extracted by mCADRE were more functionally accurate than MBA models, albeit with a long tail towards larger distances that likely reflected the bimodality of mCADRE model contents and predictions. FASTCORE models covered the narrowest range of distances and were less functionally accurate than most mCADRE models. Importantly, all MEMs outperformed the baseline defined by SALARECON, meaning that they generally produced models that were more functionally accurate than the generic template model.

**Fig. 6.**
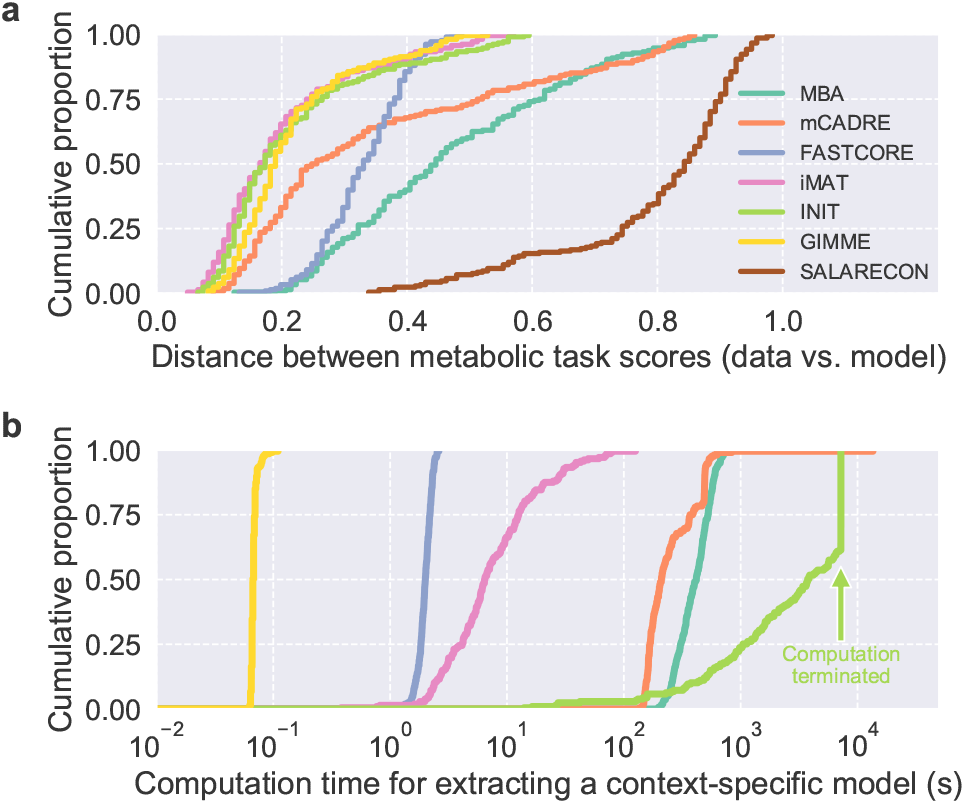
Metabolic task distance and computation time. Empirical cumulative distributions of (a) normalized Hamming distance between binary task scores inferred from data and task feasibility predicted by models, and (b) computation time required to extract models.

The iMAT-like MEMs and GIMME had very similar distributions of distances between predicted and inferred task scores, perhaps with iMAT and INIT outperforming GIMME very slightly. However, looking at distributions of computation time for model extraction, we found that GIMME was by far the most computationally efficient of all the tested MEMs (**Fig. 6**). GIMME was more than an order of magnitude faster than the other MEMs and completed nearly all extractions in less than 0.1 s. FASTCORE was also faster than most MEMs with all extractions finishing within 2.6 s. The remaining MEMs were all less computationally efficient with wider distributions of computation: from 0.4 s to 2 min for iMAT, from 3 min to 14 min for MBA, from 2 min to about 3 h 45 min for mCADRE, and from 14 s to 2 h for INIT, which was generally much slower than the other MEMs.

## Discussion

We found large variation in contents and predictions between MEMs but not between life stages or feeds, which is in line with results from systematic analyses with human models and data (Opdam *et al*., 2017; Richelle *et al*., 2019b). Supporting some of our specific findings, one study found that reaction and feasible task counts differed between MEMs largely as we describe and that MEM explained much more variance in PC scores from PCA of task feasibility than tissue or cell type (Richelle *et al*., 2019b). The only factors affecting context-specific model contents and predictions more than MEM in these studies were rules for applying constraints to the generic template model and thresholds to the transcriptomics data (Opdam *et al*., 2017; Richelle *et al*., 2019b). However, we used their recommended constraints and thresholds and therefore did not test these factors (**Table 1**). The fact that results from human studies hold also for a fish demonstrates that MEMs and their settings generalize to other animals than mammals.

MBA and mCADRE clearly differed from the other MEMs in terms of context-specific model contents and predictions. MBA models tended both to be closer to SALARECON and to have wider distributions than models built by other MEMs. This could be at least partially explained by MBA preserving two core reaction sets (high and medium confidence) rather than one, which likely leads to larger models than other MEMs. Notably, the distribution of growth rate predictions from MBA models was bimodal even though all model content distributions were unimodal. Bimodal distributions of predictions were also seen for mCADRE models, but this could be explained by bimodal distributions of contents. However, the causes of the observed bimodality in contents are unclear. Taking all MEMs together, growth rate predictions were bimodal and tended to either be close to the maximum predicted by SALARECON or close to a relative rate of about 60%, indicating that many models were reduced until only core pathways needed to produce biomass at a minimal rate remained.

FASTCORE and GIMME models preserved the growth rate and pFBA flux of SALARECON to a larger extent than other MEMs. In particular, FASTCORE preserved pFBA flux and GIMME preserved growth rate. The iMAT-like MEMs consistently produced the smallest models, which is probably why iMAT and INIT models had low growth rates, high pFBA fluxes, and low feasible task counts. Indeed, this was a tendency across all MEMs: smaller models produced biomass precursors less efficiently and performed fewer metabolic tasks. The most obvious exceptions were GIMME models, which performed fewer tasks than expected based on reaction and metabolite counts, but these counts were in turn larger than expected based on gene counts. The explanation for this is that GIMME, unlike the other MEMs, preserved all exchange reactions, which are not mapped to genes and allow metabolites to move in and out of the extracellular environment. GIMME models also preserved the growth rate and to some extent pFBA flux of SALARECON despite being substantially reduced, showing that availability of environmental metabolites to some extent can compensate for the effects of pathways lost during extraction.

There were also notable contrasts between MEMs in terms of the number of models that were able to perform metabolic tasks, reflecting the observed variation in model contents and predictions. For example, tasks in all six metabolic systems tended to be performed by the majority of MBA models, which were the largest, and by many models from the other two MBA-like MEMs – mCADRE and FASTCORE – which also produced comparatively large models. For the iMAT- and GIMME-like MEMs, most tasks were performed by a minority of models and several tasks were infeasible in all models. Perfect agreement between models on task feasibility was more common for iMAT, INIT, and GIMME than for MBA, mCADRE, and FASTCORE. It is important to note that, perhaps counterintuitively, models that perform many tasks are not necessarily better than those that perform few. A good context-specific CBM should ideally perform the tasks that are actually performed by the organism in that context, i.e., tasks for which there is evidence in literature and data.

Despite large variation in task feasibility, some tasks were frequently performed across all MEMs, and we found remarkable agreement between these tasks and metabolic processes known to be important in the liver. For example, consistent feasibility of cellular respiration and sugar degradation is in line with the liver’s role as a hub that transforms dietary nutrients into energy and building blocks for other tissues (Rui, 2014). However, energy metabolism is fundamental for any cell and its consistent feasibility could simply be driven by growth requirements in the MEMs. Other observations are less likely to be artifacts, e.g., consistent feasibility of the thioredoxin system, which can play an important role in reducing oxidative stress caused by high-fat diets (Qin *et al*., 2014), and nucleotide salvage, which is a key part of the liver’s central control of nucleotide synthesis (Fustin *et al*., 2012). The liver also synthesizes many amino acids, including glutamate, glutamine, alanine, aspartate, and glycine, which were most frequently synthesized by context-specific CBMs (Hou *et al*., 2020). Consistent feasibility of SAM synthesis reflects the fact that SAM is essential for liver health and mostly generated in hepatocytes (Mato *et al*., 2013); UDP-glucose is a precursor for glycogen, which is synthesized and stored in the fish liver (Polakof *et al*., 2012); fructose-6-phosphate synthesis through the pentose phosphate pathway is particularly important in the liver (Akram *et al*., 2019); glycerol-3-phosphate is a precursor for glycerolipids, which are predominantly synthesized in the liver (Alves-Bezerra and Cohen, 2018); and malonyl-CoA is essential for primarily hepatic fatty acid synthesis (Foster, 2012).

The liver is a very metabolically active organ, so detecting processes that occur in the liver may not be surprising. However, it is notable that the most frequently performed tasks across MEMs were known to be important for liver metabolism. This supports context-specific modeling as an approach for studying tissue-specific metabolism, and it also suggests ensemble modeling as a potential strategy for managing uncertainty and making context-specific model predictions more robust. Specifically, one could use several different MEMs and template models to build an ensemble of CBMs for each organism and context of interest, i.e. from the same omics data, and predictions could be based on agreement among models in the ensemble (Biggs and Papin, 2017). Ensemble modeling could also help improve the models themselves by applying machine learning to their contents and predictions (Medlock and Papin, 2020). Indeed, recent studies have demonstrated context-specific ensemble modeling with a single MEM (Rodríguez-Mier *et al*., 2021) and combined multiple MEMs to build a single model (Vieira *et al*., 2022).

Comparing distributions of model contents and predictions, we saw clear differences between MEMs but not between life stages or feeds. To detect contrasts between these factors among context-specific CBMs, we therefore applied PCA to reaction presence and task feasibility. Results were very similar for reactions and tasks, reflecting the fact that task feasibility is determined by reaction presence. In both cases, the first five PCs explained about half of the total variance in the data, and variance in the scores of these PCs was mainly explained by MEM. Life stage or feed explained comparatively tiny amounts of variance, but hierarchical clustering revealed a tendency for grouping by life stage within MEMs, leading us to perform PCA separately for each MEM. This did indeed lead to PCs capturing differences between life stages, most notably for GIMME, FASTCORE, and MBA, but not between feeds. Performing another set of PCAs within each MEM and life stage also failed to separate models based on diet, possibly due to the simplified representation of lipids in SALARECON. Future studies should expand lipid metabolism in SALARECON to potentially detect differences between feeds. These results show that choice of MEM is by far the most important determinant for model contents and predictions, but at least some of the MEMs are capable of producing models that capture biological differences.

Finally, we evaluated the functional accuracy and computational efficiency of MEMs. We compared metabolic task scores inferred from the data to scores predicted by context-specific models and found that agreement between inferred and predicted scores tended to be the highest for iMAT, INIT, and GIMME. These were also the MEMs that produced the smallest models, meaning that smaller models tended to be more context-specific than larger ones. FASTCORE models were the most consistent, although they performed slightly worse the iMAT- and GIMME-like models. MBA and mCADRE both had long tails, and some models performed as poorly as SALARECON, the generic template model. GIMME was an order of magnitude faster than FASTCORE, which was in turn faster than iMAT. MBA and mCADRE both required about equally long computation times, while INIT was the slowest, requiring us to terminate the procedure after two hours. A near-optimal model was always found within this time in a previous study (Opdam *et al*., 2017). The differences in efficiency between MEMs are largely as expected based on their optimization strategies: GIMME solves a fixed number of linear programs (LPs), while other MEMs either solve more LPs or much harder mixed-integer linear programs (Robaina Estévez and Nikoloski, 2014).

There are many other MEMs available that were not systematically tested in this or other studies, including recent methods that account for trancriptomic variability (Joshi *et al*., 2020), use ensemble modeling to improve predictions (Rodríguez-Mier *et al*., 2021), or combine multiple methods and settings (Vieira *et al*., 2022). Notably, some methods integrate metabolic tasks into the model extraction procedure itself by requiring agreement with inferred task feasibility for human models and data (Agren *et al*., 2014; Richelle *et al*., 2019b). This can increase consensus among context-specific CBMs aross MEMs (Richelle *et al*., 2019b) but it does not necessarily improve model contents and predictions relative to simpler methods such as GIMME (Lee *et al*., 2022). There have also been numerous efforts to develop MEMs that integrate multiple types of omics data (Cho *et al*., 2019). In this study, we only focused on transcriptomics data, which is the exclusively accepted data type of most MEMs (Cho *et al*., 2019), and we used metabolic tasks to evaluate the performance of MEMs that have been systematically tested for human applications (Robaina Estévez and Nikoloski, 2014; Opdam *et al*., 2017; Richelle *et al*., 2019b). We found that several of the tested MEMs captured expected task feasibility well without enforcing it in the procedure, showing that simple MEMs informed only by transcriptomics can capture key biological differences between contexts. Moreover, all of these MEMs can be extended to integrate tasks (Richelle *et al*., 2019a) and multi-omics data (Cho *et al*., 2019), so knowing which MEMs likely provide the best baseline will be useful for future studies of context-specific Atlantic salmon metabolism.

## 4 Conclusion

Contents and predictions of context-specific CBMs were mainly determined by MEM, but life stage explained significant variance in PC scores for reactions (GIMME, FASTCORE, MBA, and iMAT) and tasks (GIMME and FASTCORE). The iMAT- and GIMME-like MEMs – iMAT, INIT, and GIMME – captured context-specific metabolic activities inferred from the data better than the other MEMs, and GIMME was by far the fastest method. All things considered, GIMME seems to have a slight advantage over other MEMs; it is simple but fast and produces models that are remarkably consistent in their contents and predictions as well as functionally accurate. Compared to GIMME, FASTCORE captured life stages equally well but was slower and less functionally accurate, while iMAT and INIT captured life stages less well and were slower but equally functionally accurate. Context-specific CBMs consistently outperformed SALARECON and metabolic tasks predicted to be feasible across all MEMs were remarkably liver-specific, showing that context-specific modeling can improve predictions and explain omics data available for Atlantic salmon (Samy *et al*., 2017; Yang *et al*., 2020).

## Acknowledgements

We thank Yang Jin, Hanna Sahlström, and Ylva Wedmark for helpful discussions and feedback on the manuscript.

---

**Fig. S1.**
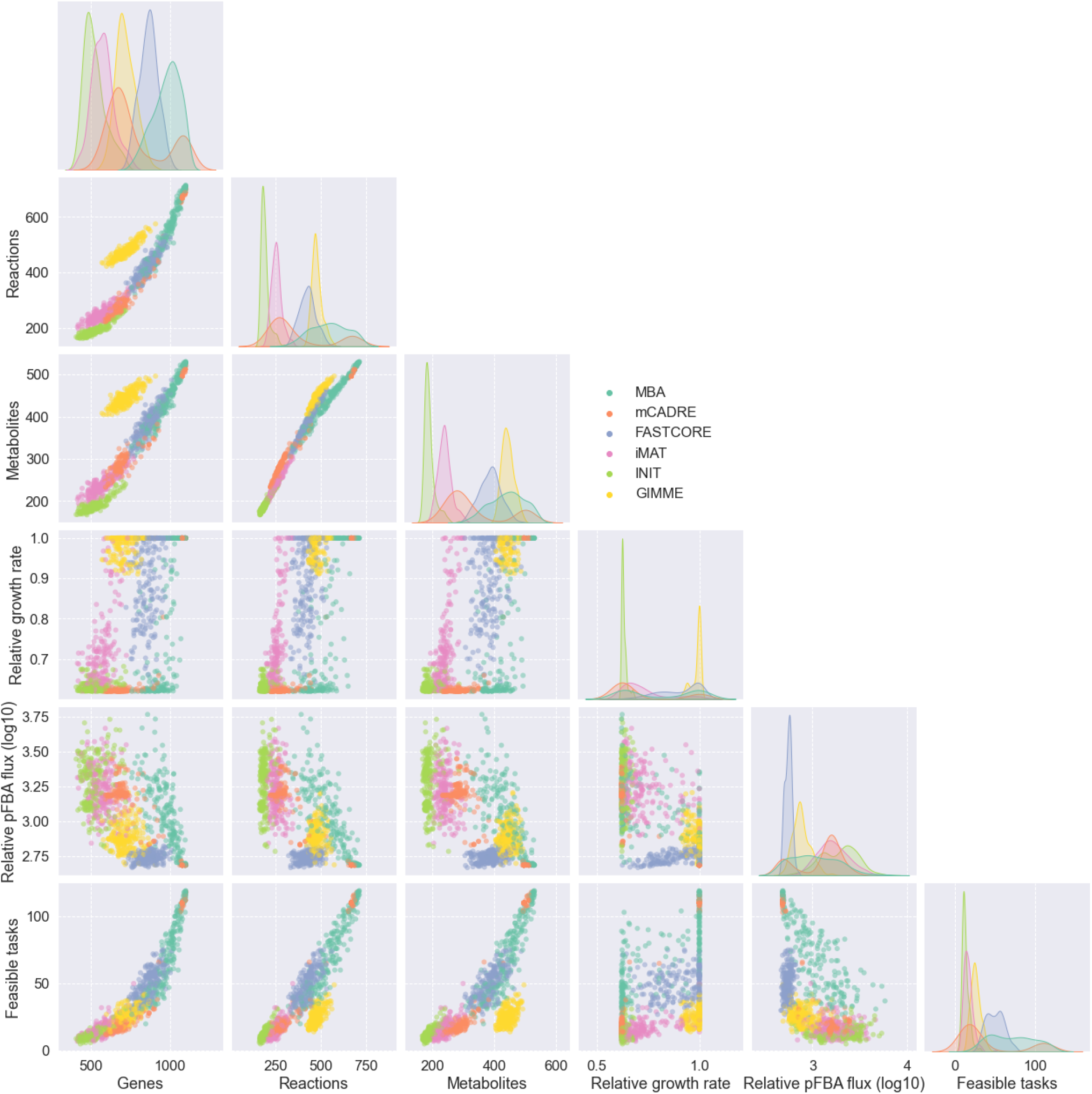
Pairwise relationships between context-specific model contents and predictions by MEM. Gene counts, reaction counts, metabolite counts, predicted maximal growth rate relative to SALARECON, sum of absolute fluxes from pFBA relative to growth rate, and feasible metabolic task counts are shown.

**Fig. S2.**
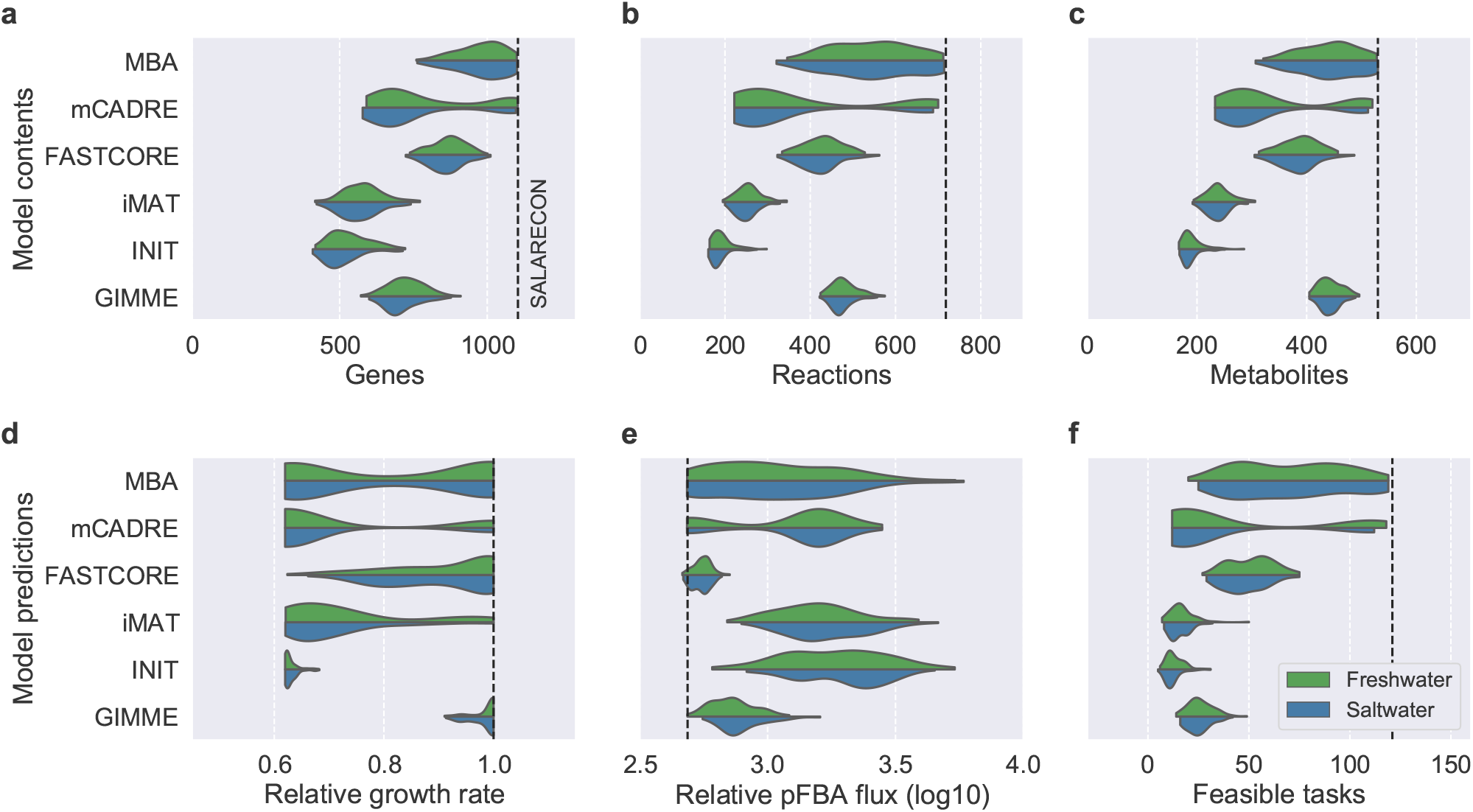
Distribution of context-specific model contents (a–c) and predictions (d–f) by MEM and life stage. (a) Gene counts, (b) reaction counts, (c) metabolite counts, (d) predicted maximal growth rate relative to SALARECON, (e) sum of absolute fluxes from pFBA relative to growth rate, and (f) feasible metabolic task counts. Kernel density estimates are scaled to the same width with cutoffs at the extreme data points. Dashed lines indicate predictions from SALARECON.

**Fig. S3.**
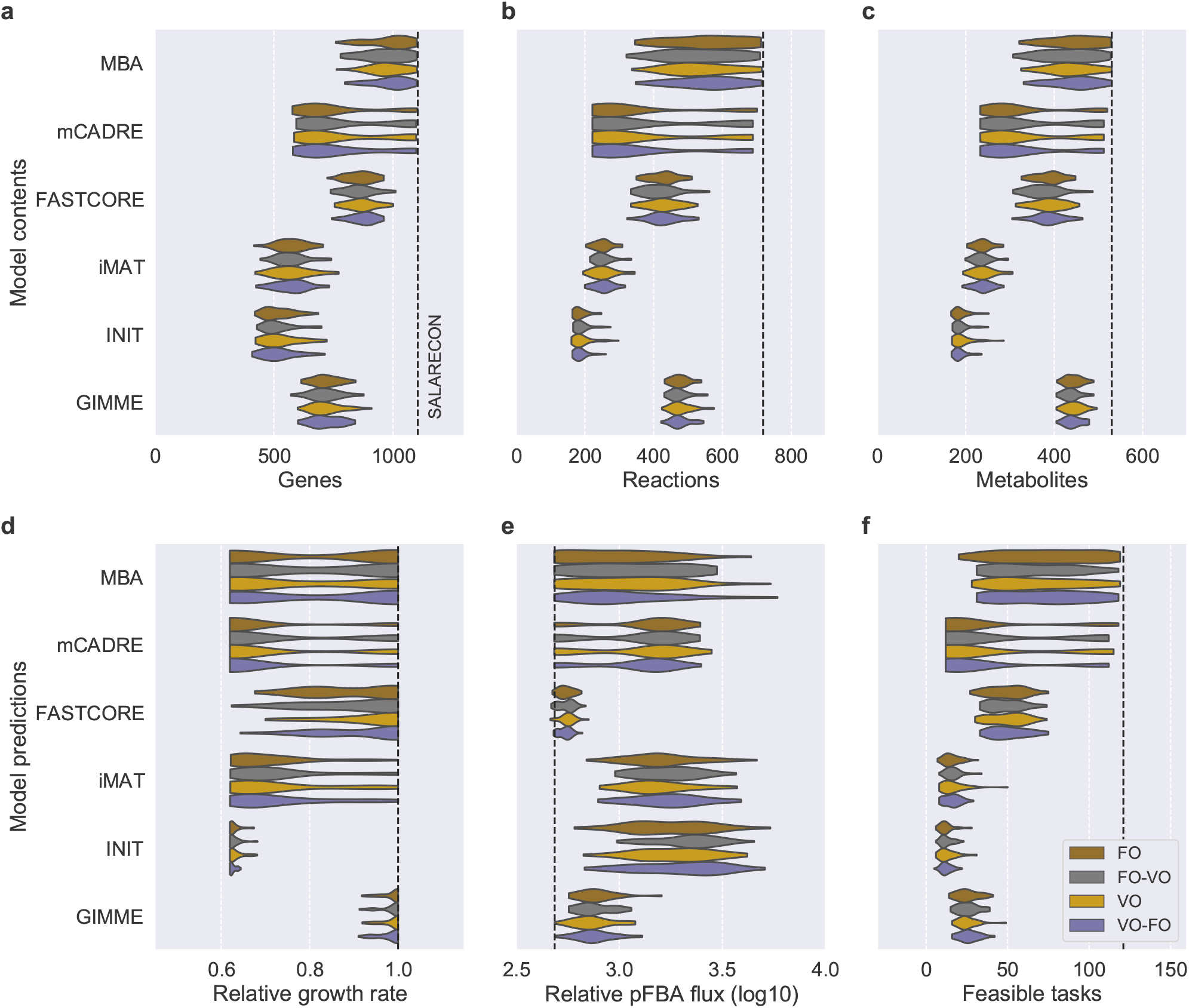
Contents and predictions of context-specific models by diet. Distribution of context-specific model contents (a–c) and predictions (d–f) by MEM and diet. (a) Gene counts, (b) reaction counts, (c) metabolite counts, (d) predicted maximal growth rate relative to SALARECON, (e) sum of absolute fluxes from pFBA relative to growth rate, and (f) feasible metabolic task counts. Kernel density estimates are scaled to the same width with cutoffs at the extreme data points. Dashed lines indicate predictions from SALARECON.

**Fig. S4.**
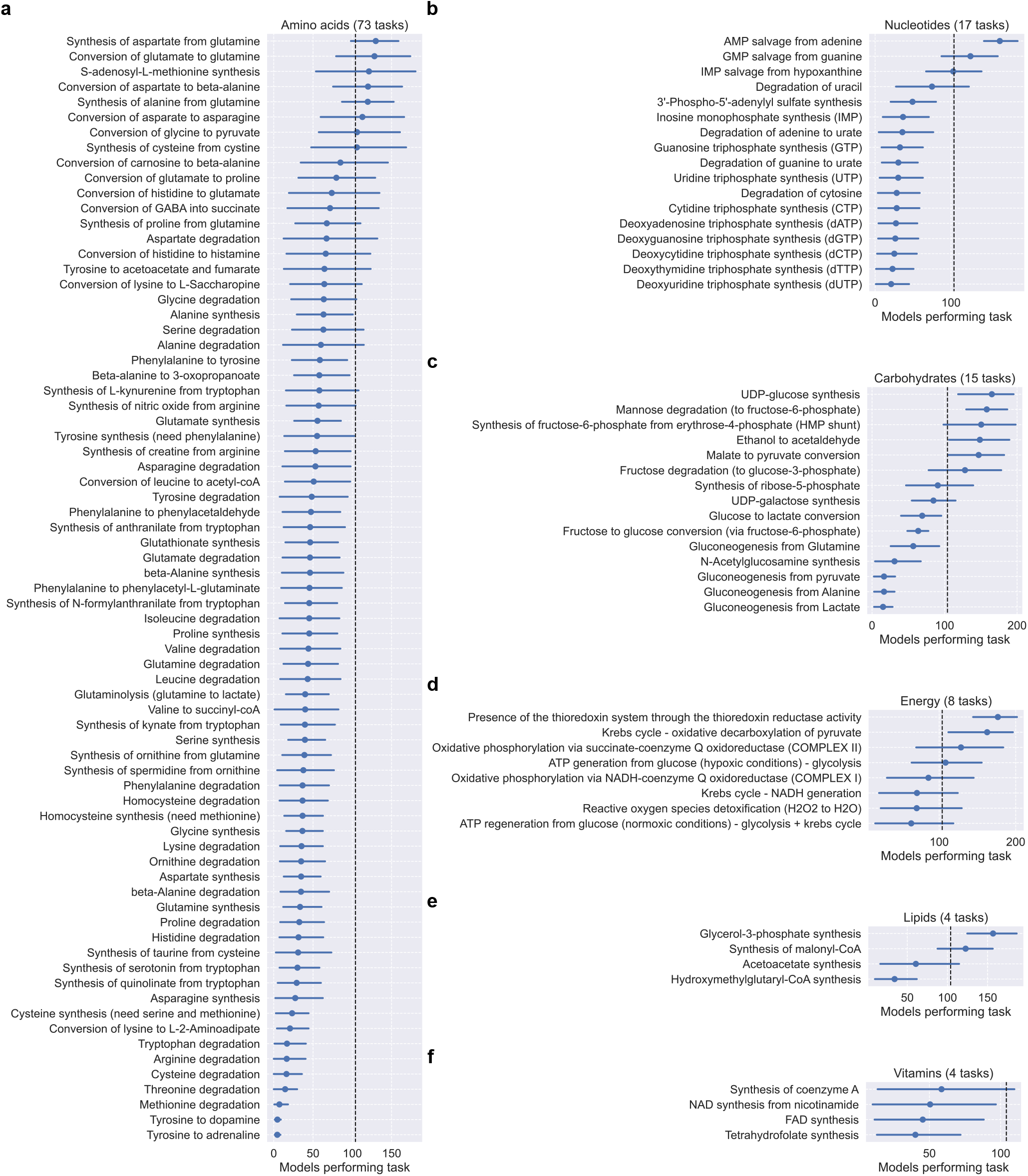
Mean number of context-specific models performing each metabolic task Mean number of context-specific models in which each metabolic task is feasible across MEMs. Tasks are divided into six metabolic systems: (a) amino acid, (b) nucleotide, (c) carbohydrate, (d) energy, (e) lipid, and (f) vitamin metabolism. Error bars indicate 95% confidence intervals for the estimated means obtained from bootstrapping with 1,000 samples. Dashed lines indicate half the number of models extracted with each MEM.

**Fig. S5.**
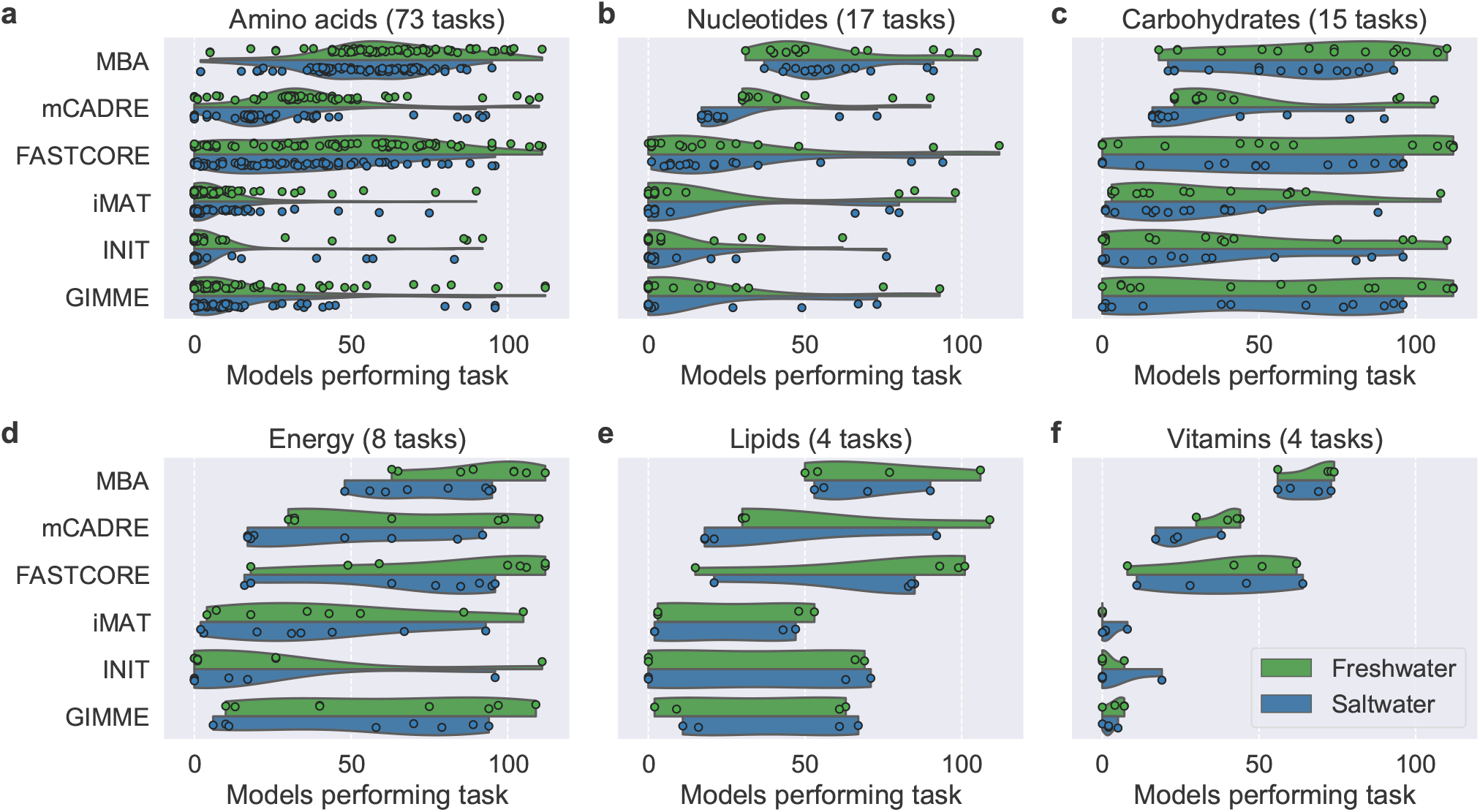
Number of context-specific models in which metabolic tasks are feasible by MEM and life stage. Markers represent tasks and are divided into six metabolic systems: (a) amino acid, (b) nucleotide, (c) carbohydrate, (d) energy, (e) lipid, and (f) vitamin metabolism. Kernel density estimates are scaled to the same width with cutoffs at the extreme data points.

**Fig. S6.**
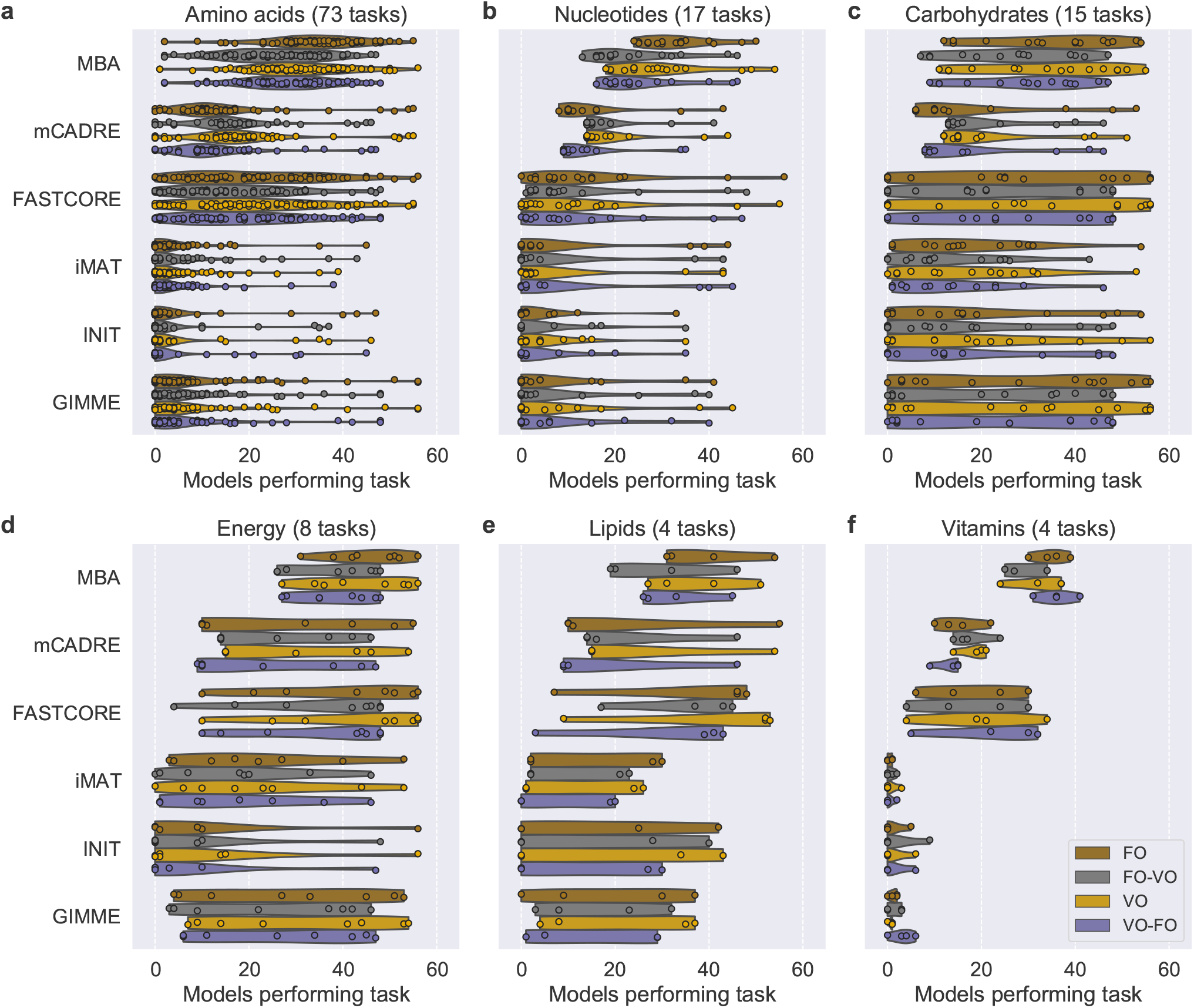
Number of context-specific models in which metabolic tasks are feasible by MEM and feed. Markers represent tasks and are divided into six metabolic systems: (a) amino acid, (b) nucleotide, (c) carbohydrate, (d) energy, (e) lipid, and (f) vitamin metabolism. Kernel density estimates are scaled to the same width with cutoffs at the extreme data points.

**Fig. S7.**
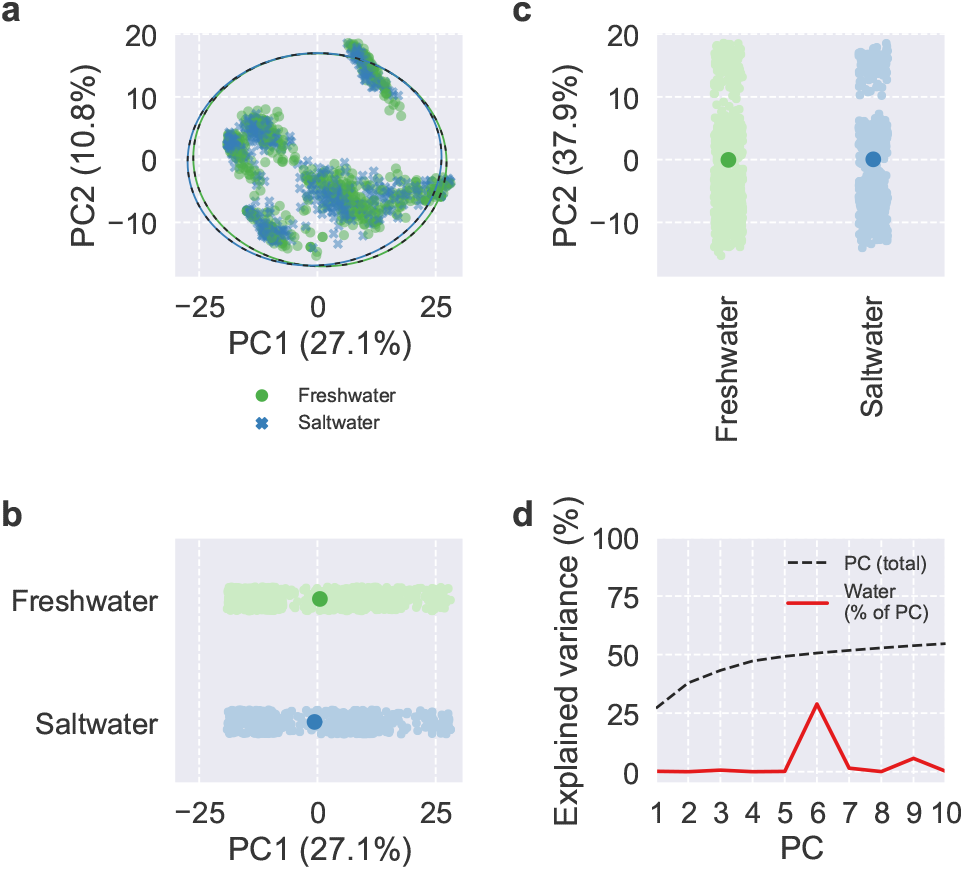
PCA of reaction presence. Scores and explained variance from PCA of reaction presence. (a–c) Scores of the first two PCs, colored by life stage, with 95% confidence ellipses and intervals. (d) Cumulative total variance explained by the first ten PCs and variance of PC scores explained by life stage.

**Fig. S8.**
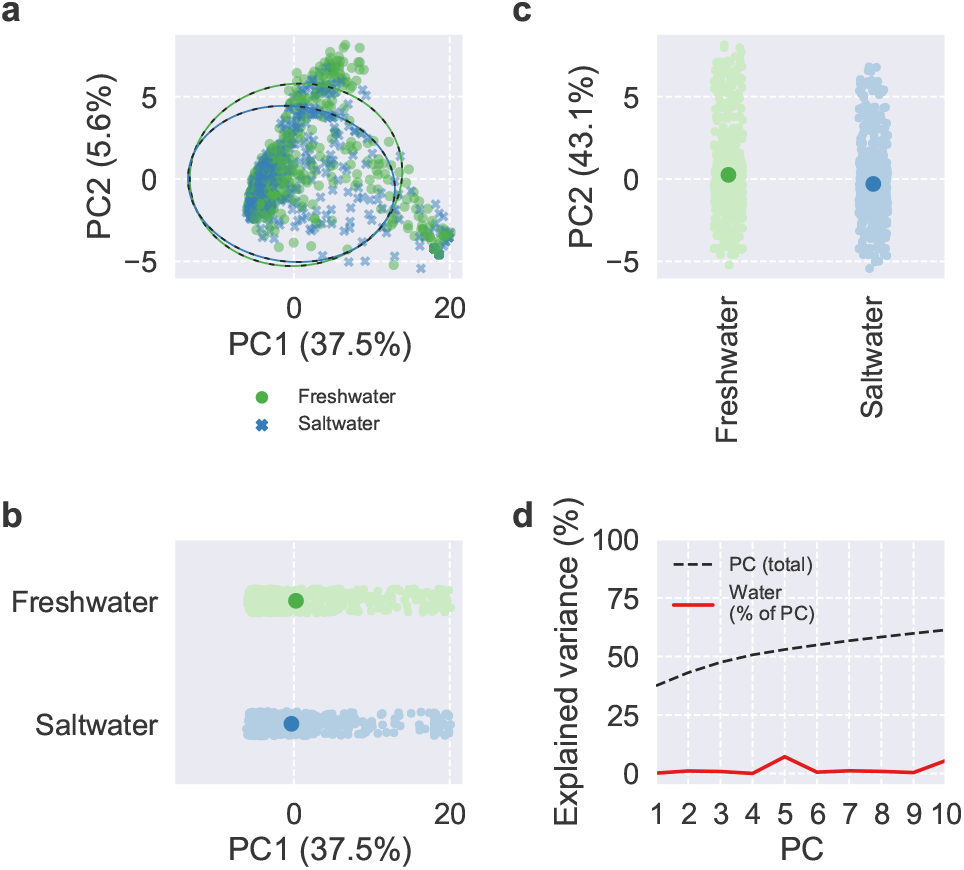
PCA of task feasibility. Scores and explained variance from PCA of task feasibility. (a–c) Scores of the first two PCs, colored by life stage, with 95% confidence ellipses and intervals. (d) Cumulative total variance explained by the first ten PCs and variance of PC scores explained by life stage.

**Fig. S9.**
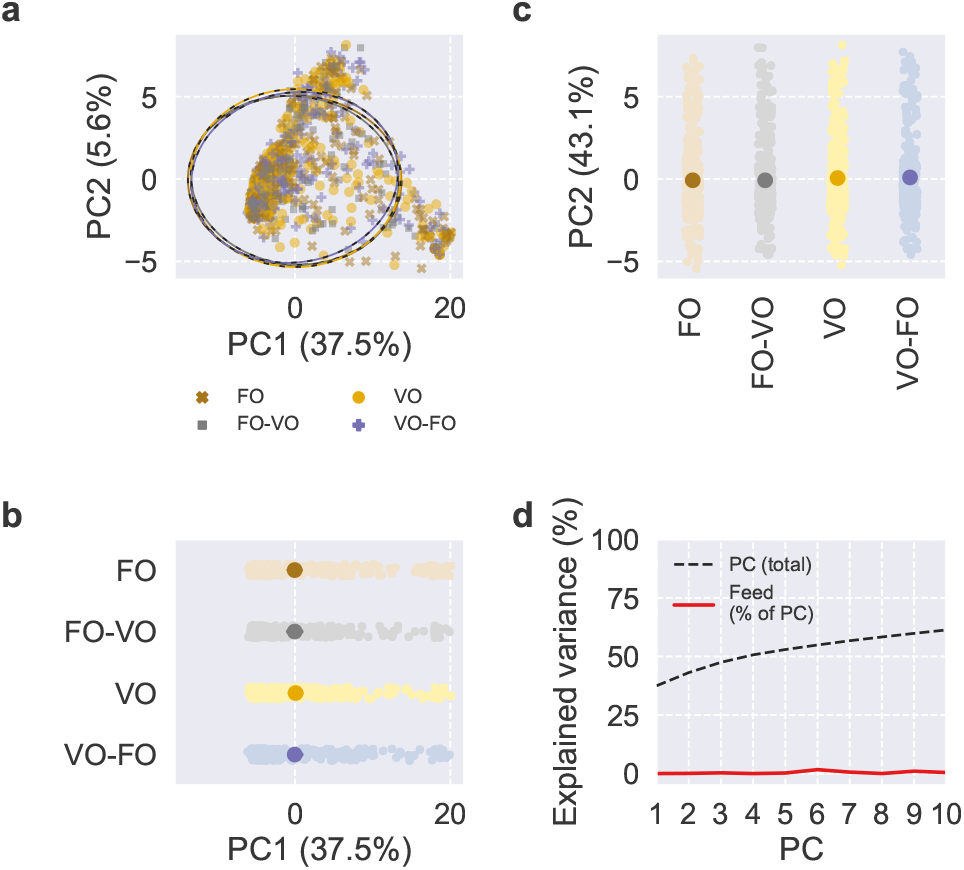
PCA of reaction presence. Scores and explained variance from PCA of reaction presence. (a–c) Scores of the first two PCs, colored by feed, with 95% confidence ellipses and intervals. (d) Cumulative total variance explained by the first ten PCs and variance of PC scores explained by feed.

**Fig. S10.**
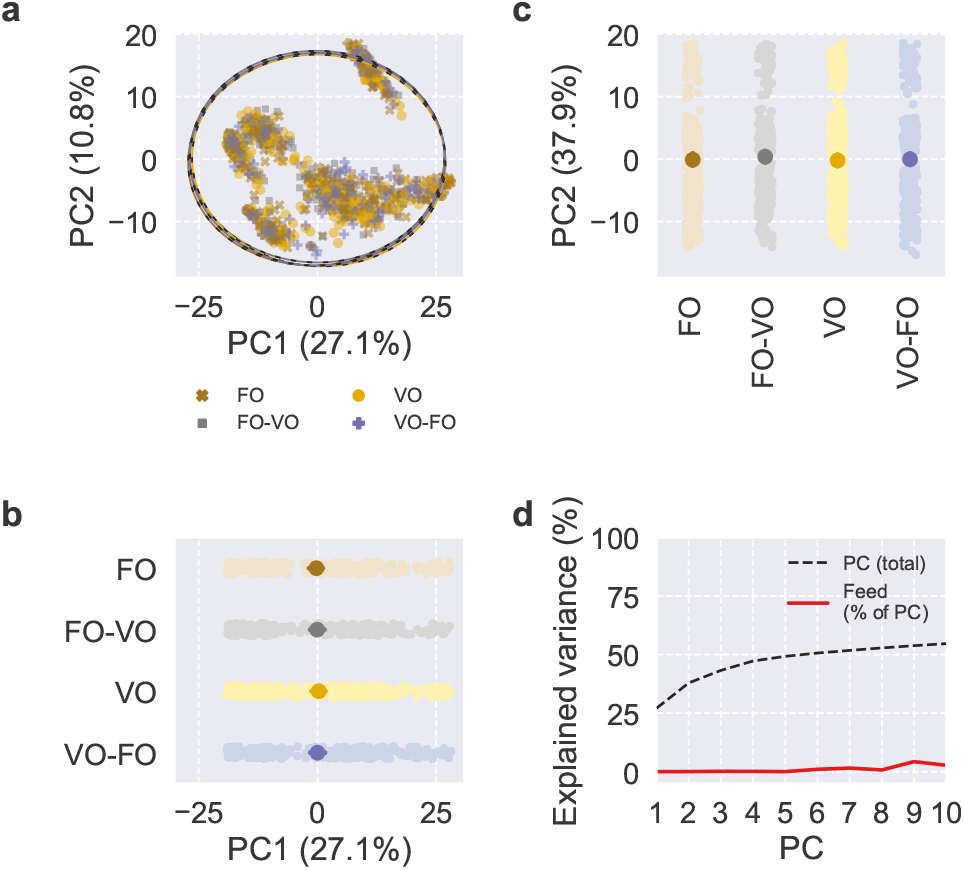
PCA of task feasibility. Scores and explained variance from PCA of task feasibility. (a–c) Scores of the first two PCs, colored by feed, with 95% confidence ellipses and intervals. (d) Cumulative total variance explained by the first ten PCs and variance of PC scores explained by feed.

**Fig. S11.**
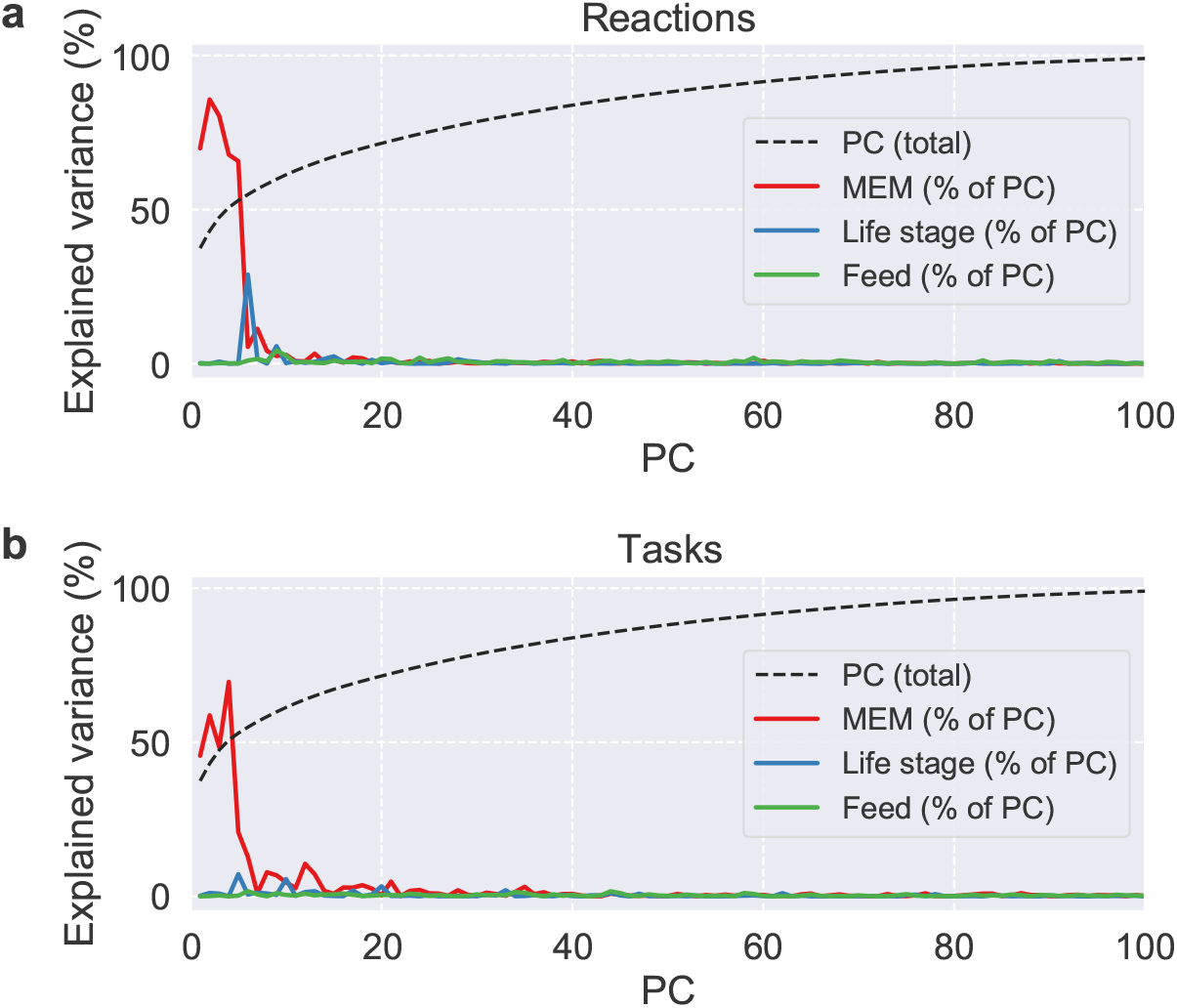
Cumulative total variance explained by the first 100 PCs and variance of PC scores explained by MEM, life stage, and feed from PCA of reaction presence and task feasibility.

**Fig. S12.**
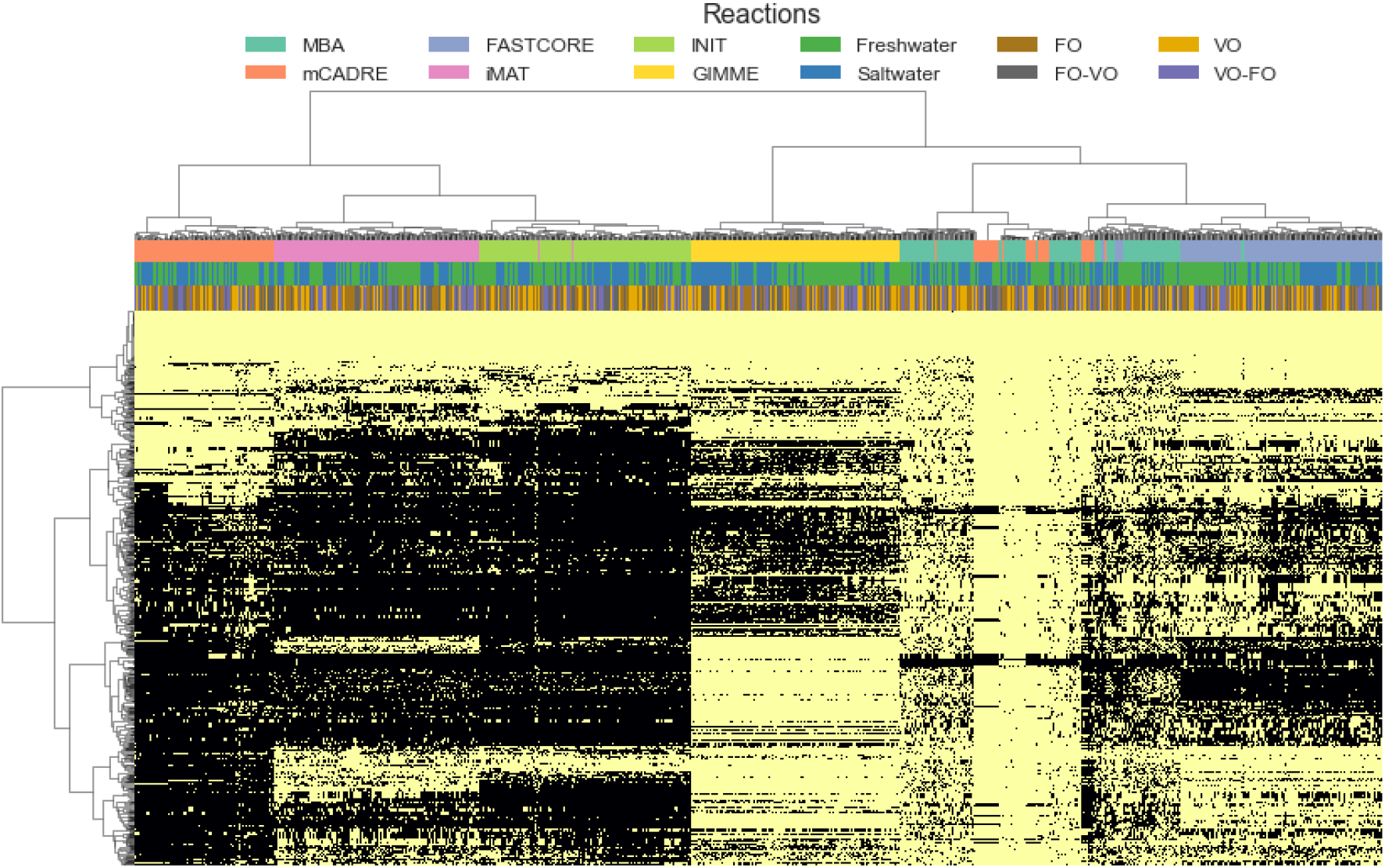
Clustered heatmap of reaction presence. Rows are reactions, columns are models (samples), and a yellow cell indicates that a reaction is present in a model. Rows and columns are clustered by Ward’s minimum variance method and columns are colored by MEM, life stage, and diet.

**Fig. S13.**
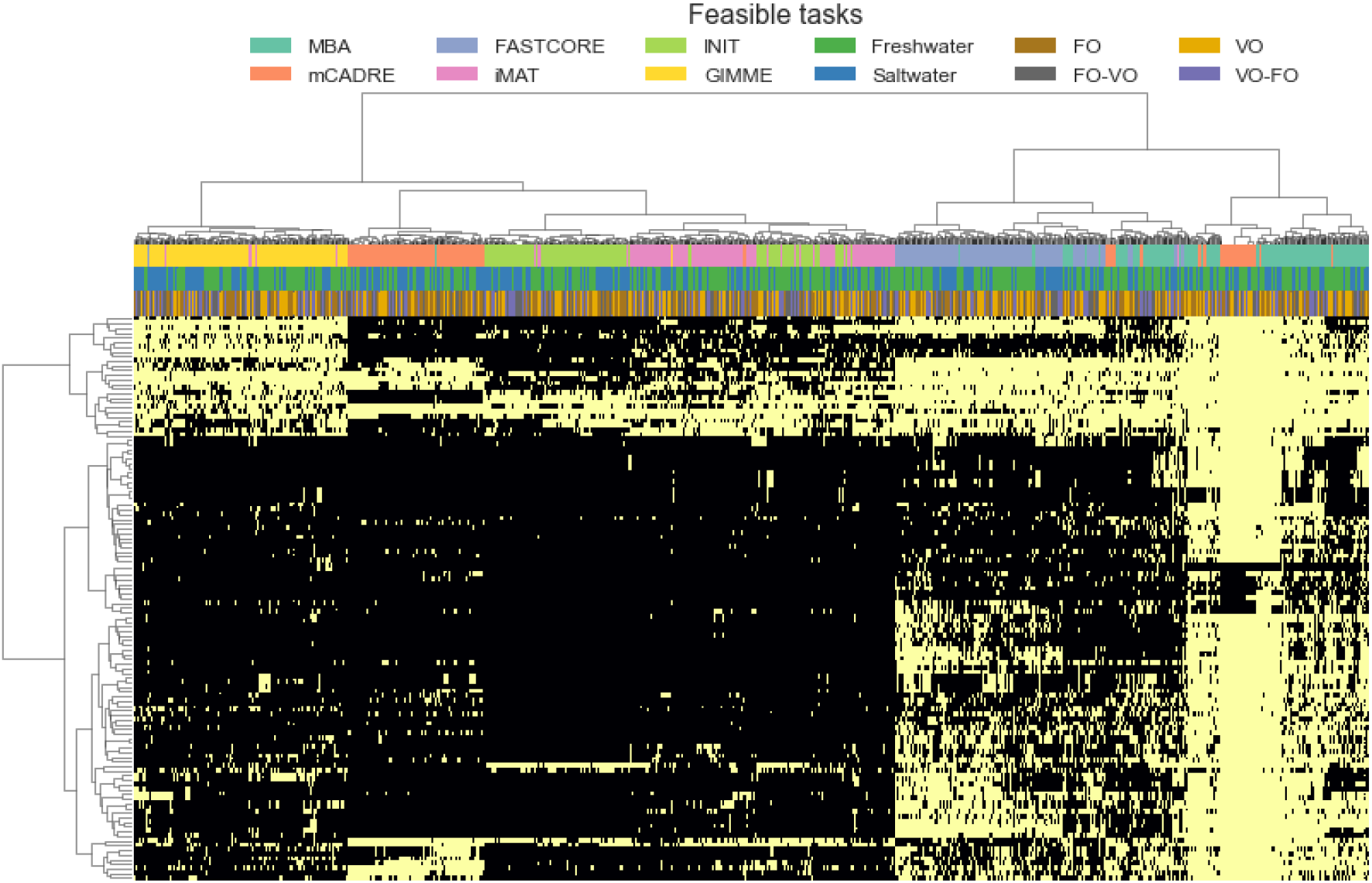
Clustered heatmap of task feasibility. Rows are metabolic tasks, columns are models (samples), and a yellow cell indicates that a task can be performed by a model. Rows and columns are clustered by Ward’s minimum variance method and columns are colored by MEM, life stage, and diet.

**Fig. S14.**
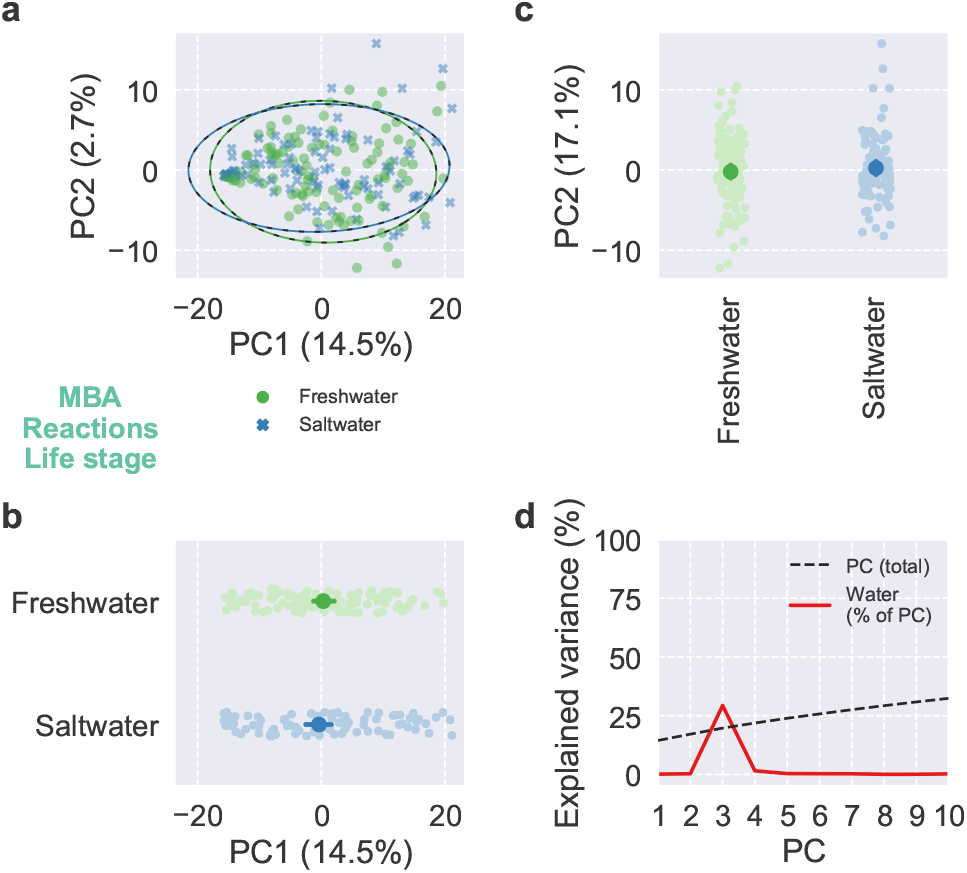
PCA of reaction presence for MBA. Scores and explained variance from PCA of reaction presence for MBA. (a–c) Scores of the first two PCs, colored by life stage, with 95% confidence ellipses and intervals. (d) Cumulative total variance explained by the first ten PCs and variance of PC scores explained by life stage.

**Fig. S15.**
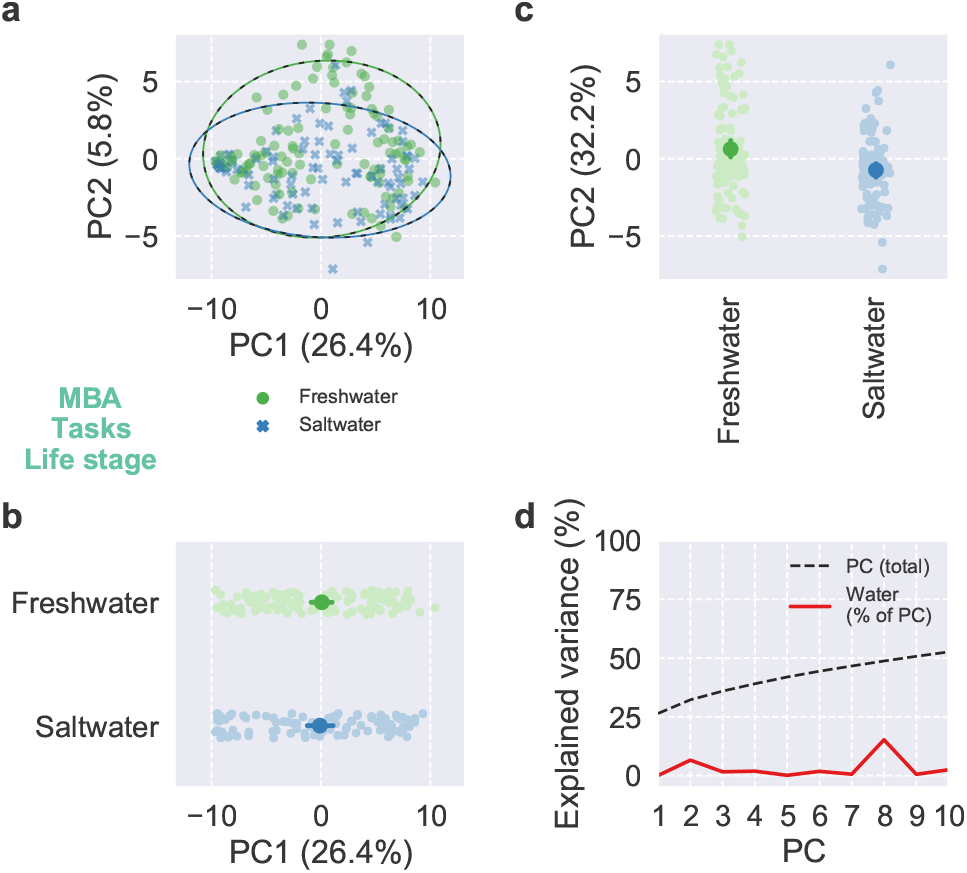
PCA of task feasibility for MBA. Scores and explained variance from PCA of task feasibility for MBA. (a–c) Scores of the first two PCs, colored by life stage, with 95% confidence ellipses and intervals. (d) Cumulative total variance explained by the first ten PCs and variance of PC scores explained by life stage.

**Fig. S16.**
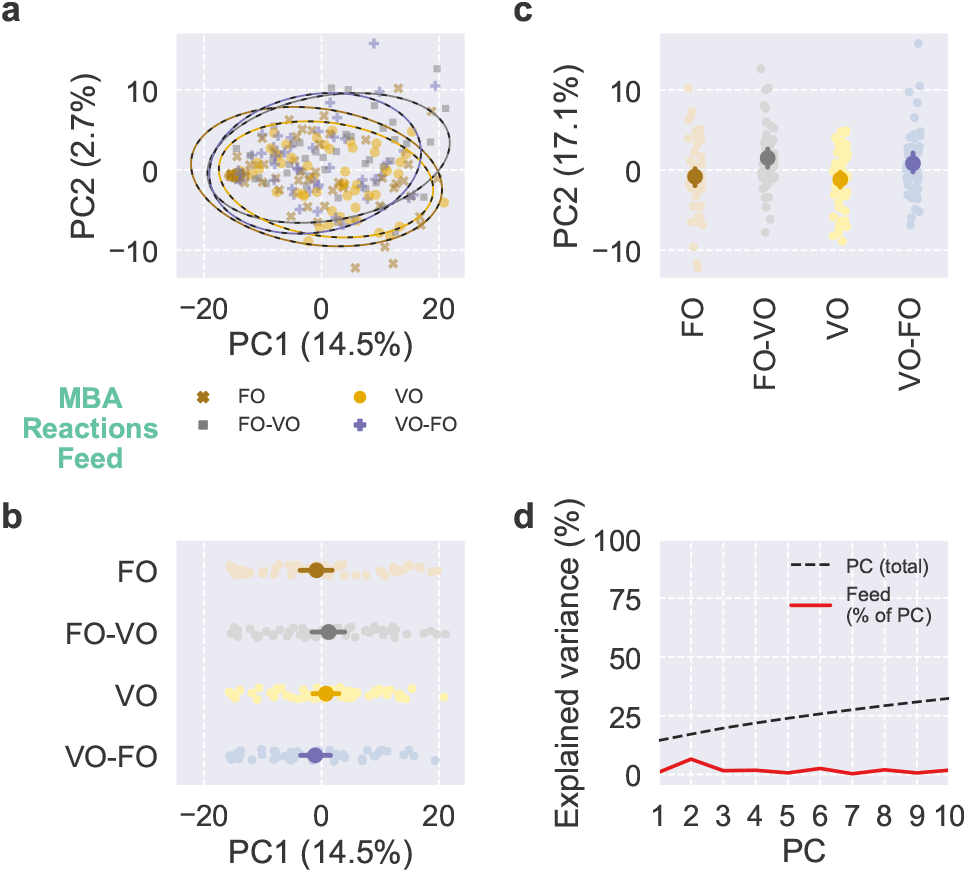
PCA of reaction presence for MBA. Scores and explained variance from PCA of reaction presence for MBA. (a–c) Scores of the first two PCs, colored by feed, with 95% confidence ellipses and intervals. (d) Cumulative total variance explained by the first ten PCs and variance of PC scores explained by feed.

**Fig. S17.**
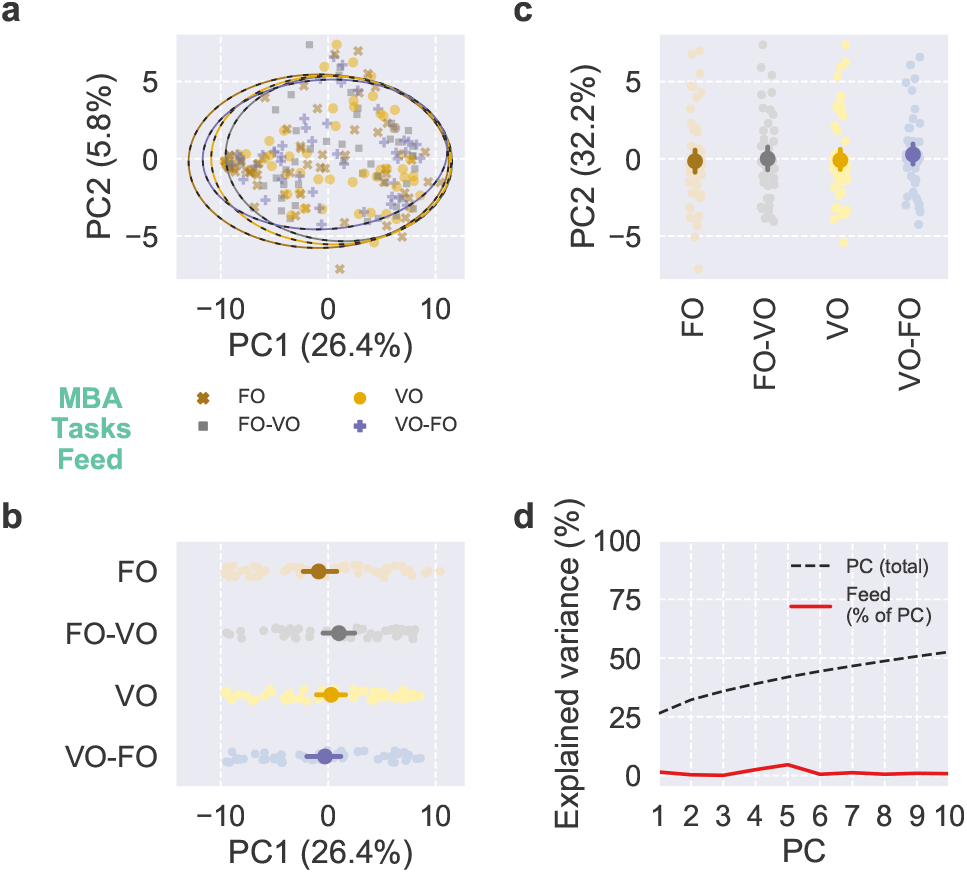
PCA of task feasibility for MBA. Scores and explained variance from PCA of task feasibility for MBA. (a–c) Scores of the first two PCs, colored by feed, with 95% confidence ellipses and intervals. (d) Cumulative total variance explained by the first ten PCs and variance of PC scores explained by feed.

**Fig. S18.**
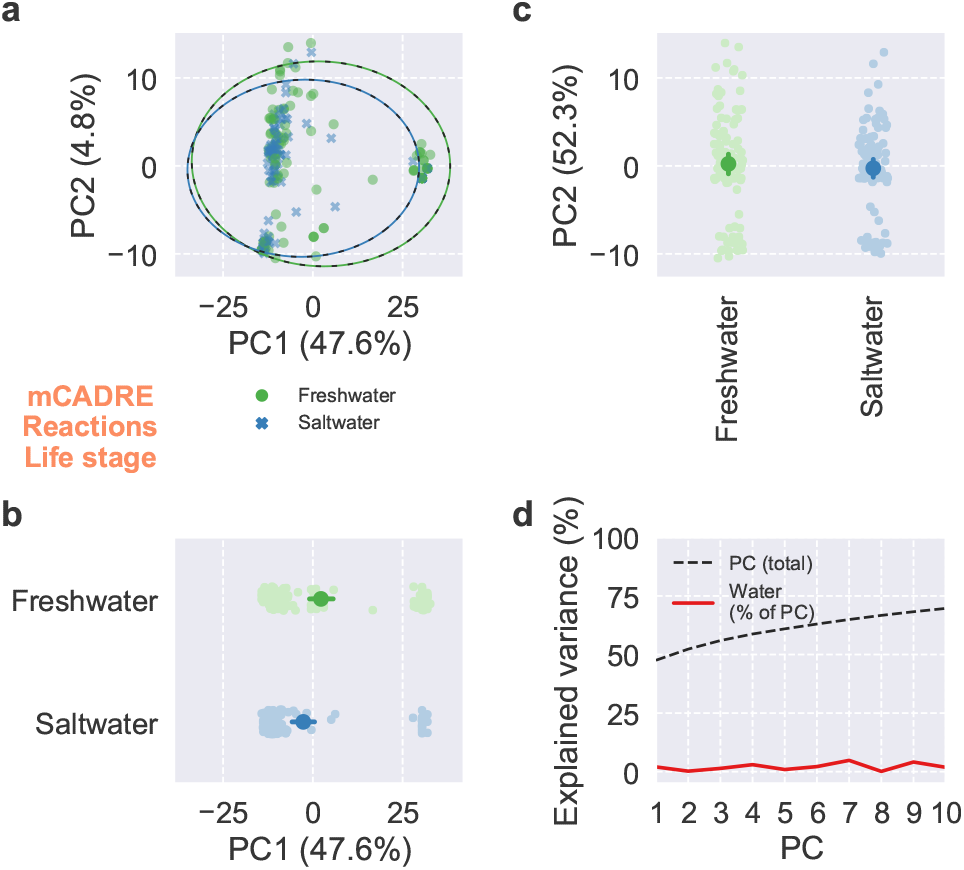
PCA of reaction presence for mCADRE. Scores and explained variance from PCA of reaction presence for mCADRE. (a–c) Scores of the first two PCs, colored by life stage, with 95% confidence ellipses and intervals. (d) Cumulative total variance explained by the first ten PCs and variance of PC scores explained by life stage.

**Fig. S19.**
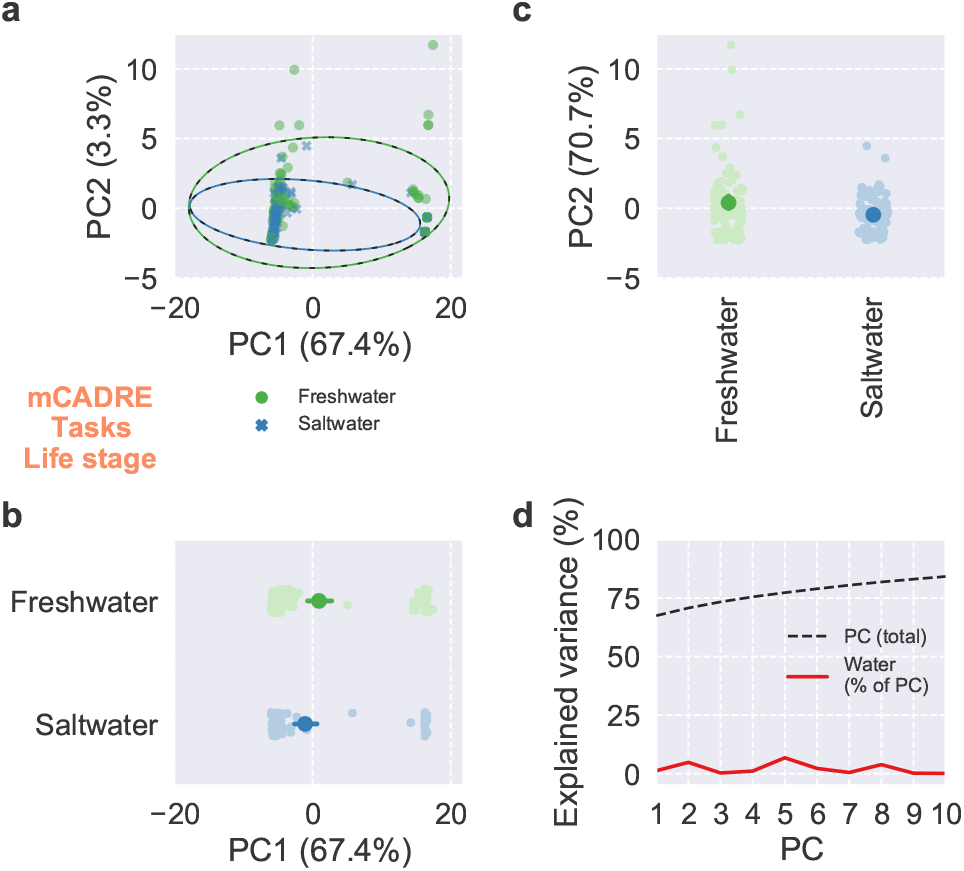
PCA of task feasibility for mCADRE. Scores and explained variance from PCA of task feasibility for mCADRE. (a–c) Scores of the first two PCs, colored by life stage, with 95% confidence ellipses and intervals. (d) Cumulative total variance explained by the first ten PCs and variance of PC scores explained by life stage.

**Fig. S20.**
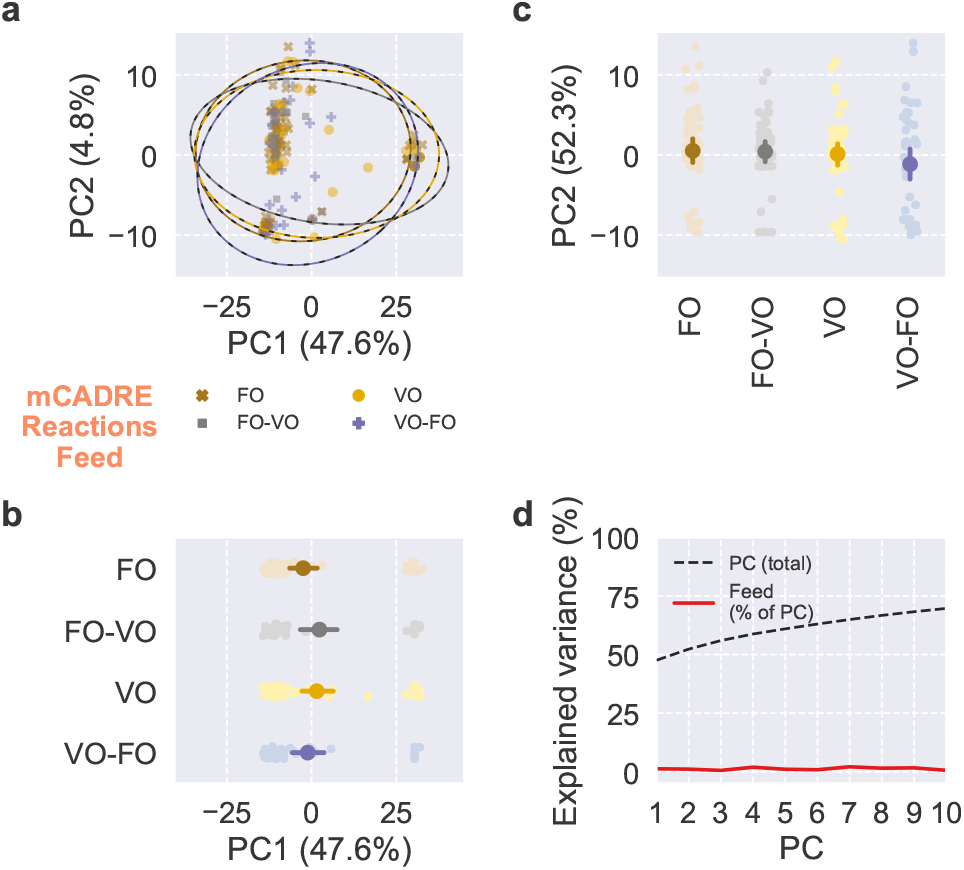
PCA of reaction presence for mCADRE. Scores and explained variance from PCA of reaction presence for mCADRE. (a–c) Scores of the first two PCs, colored by feed, with 95% confidence ellipses and intervals. (d) Cumulative total variance explained by the first ten PCs and variance of PC scores explained by feed.

**Fig. S21.**
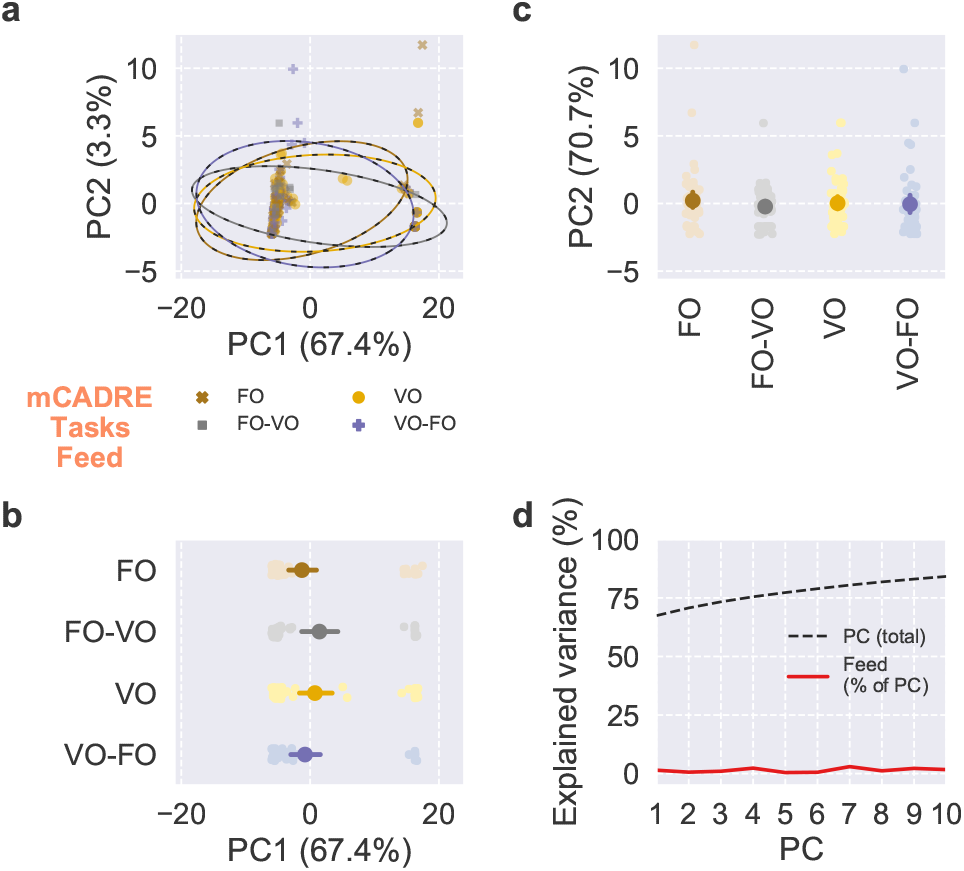
PCA of task feasibility for mCADRE. Scores and explained variance from PCA of task feasibility for mCADRE. (a–c) Scores of the first two PCs, colored by feed, with 95% confidence ellipses and intervals. (d) Cumulative total variance explained by the first ten PCs and variance of PC scores explained by feed.

**Fig. S22.**
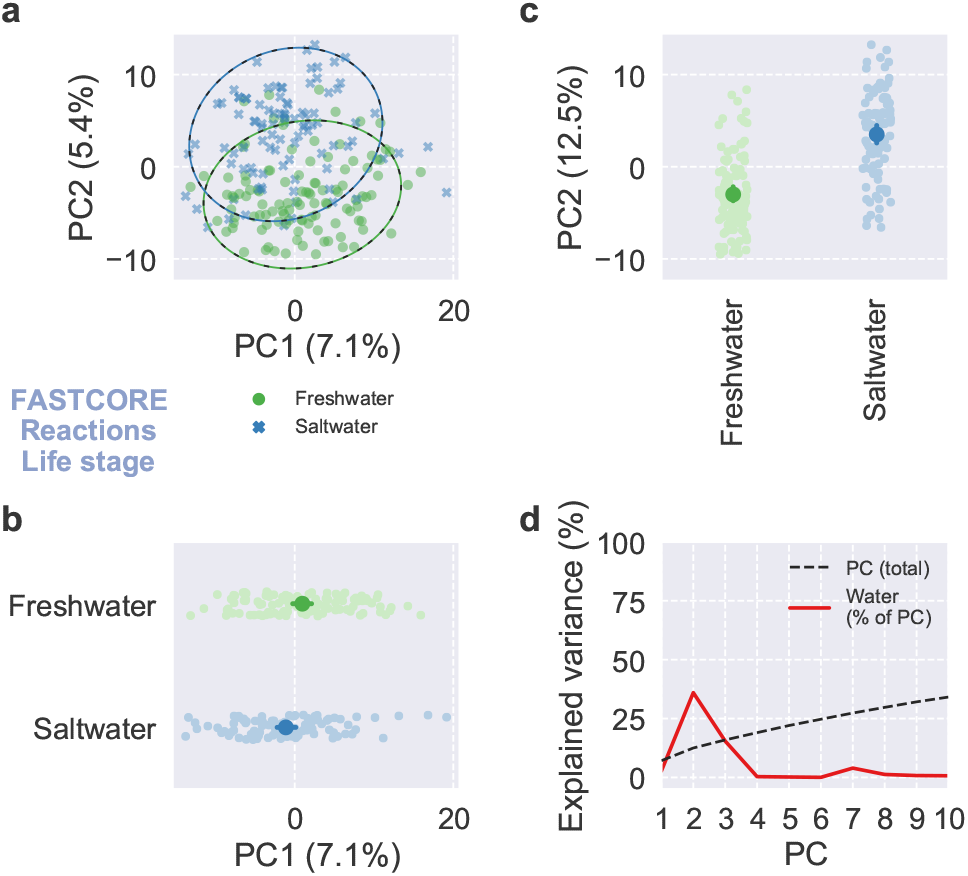
PCA of reaction presence for FASTCORE. Scores and explained variance from PCA of reaction presence for FASTCORE. (a–c) Scores of the first two PCs, colored by life stage, with 95% confidence ellipses and intervals. (d) Cumulative total variance explained by the first ten PCs and variance of PC scores explained by life stage.

**Fig. S23.**
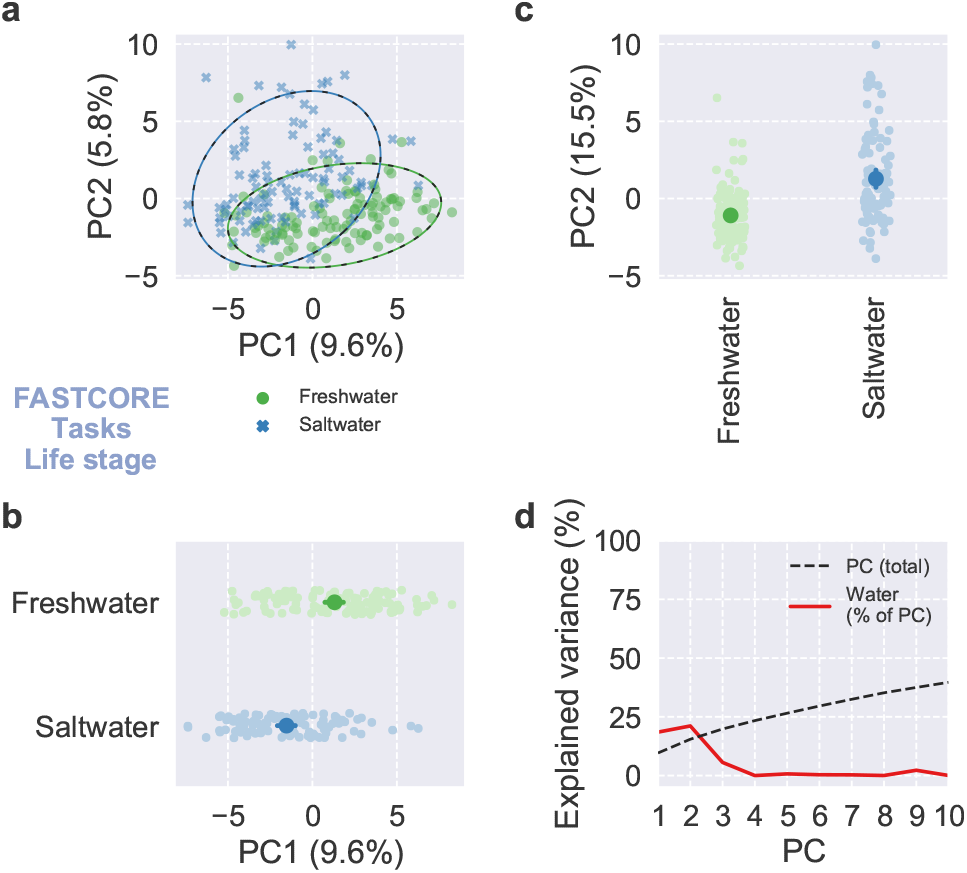
PCA of task feasibility for FASTCORE. Scores and explained variance from PCA of task feasibility for FASTCORE. (a–c) Scores of the first two PCs, colored by life stage, with 95% confidence ellipses and intervals. (d) Cumulative total variance explained by the first ten PCs and variance of PC scores explained by life stage.

**Fig. S24.**
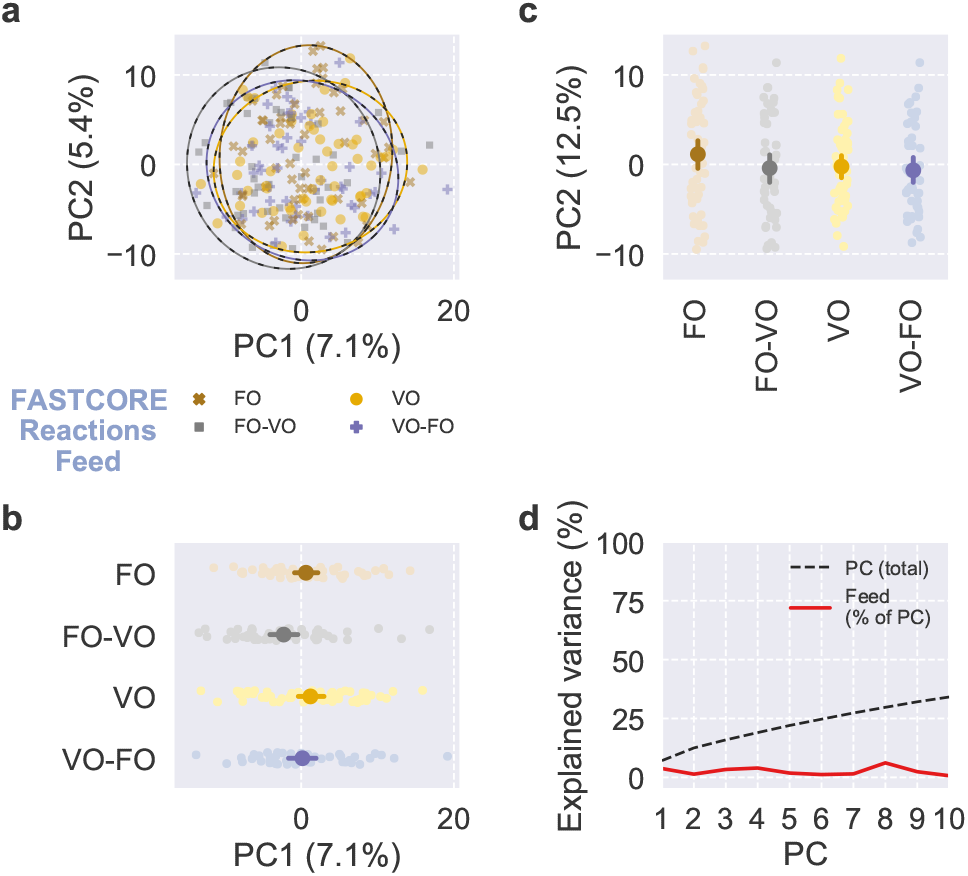
PCA of reaction presence for FASTCORE. Scores and explained variance from PCA of reaction presence for FASTCORE. (a–c) Scores of the first two PCs, colored by feed, with 95% confidence ellipses and intervals. (d) Cumulative total variance explained by the first ten PCs and variance of PC scores explained by feed.

**Fig. S25.**
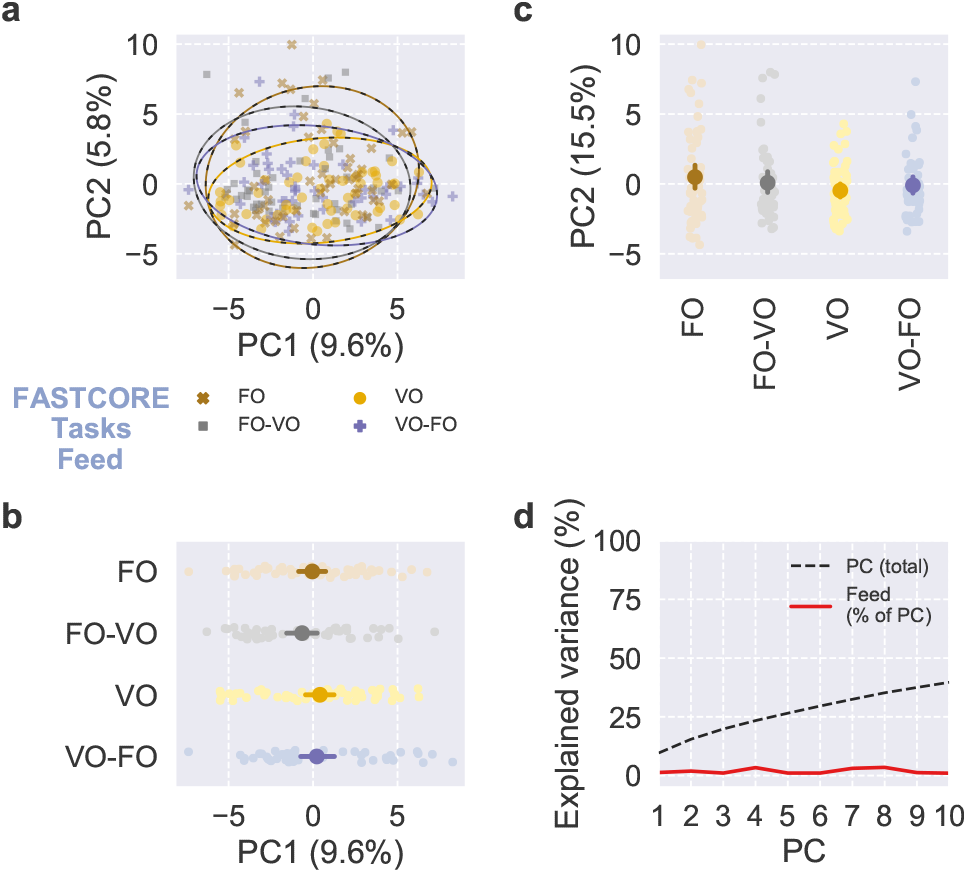
PCA of task feasibility for FASTCORE. Scores and explained variance from PCA of task feasibility for FASTCORE. (a–c) Scores of the first two PCs, colored by feed, with 95% confidence ellipses and intervals. (d) Cumulative total variance explained by the first ten PCs and variance of PC scores explained by feed.

**Fig. S26.**
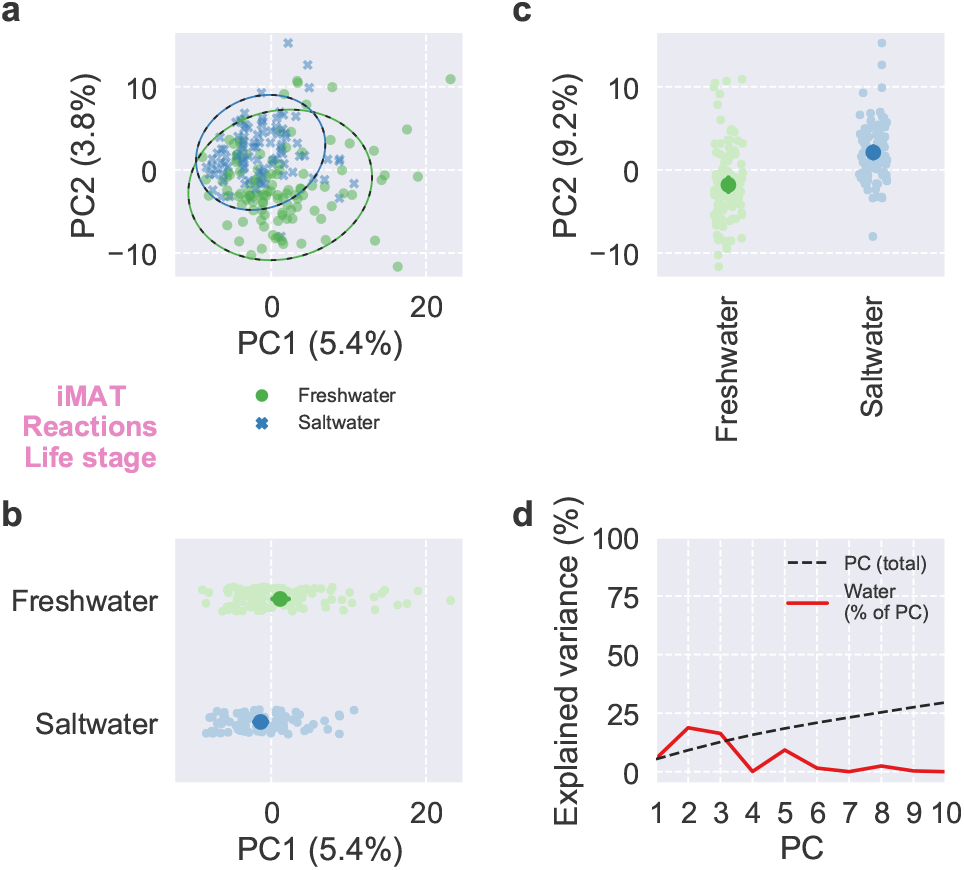
PCA of reaction presence for iMAT. Scores and explained variance from PCA of reaction presence for iMAT. (a–c) Scores of the first two PCs, colored by life stage, with 95% confidence ellipses and intervals. (d) Cumulative total variance explained by the first ten PCs and variance of PC scores explained by life stage.

**Fig. S27.**
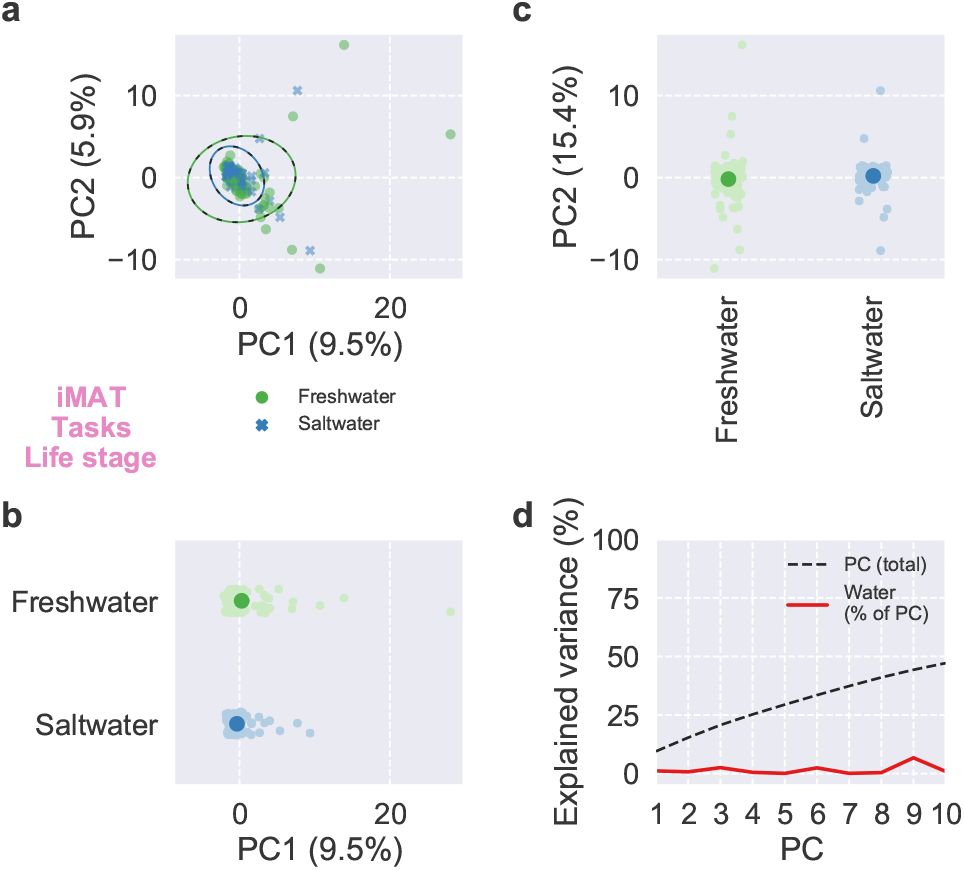
PCA of task feasibility for iMAT. Scores and explained variance from PCA of task feasibility for iMAT. (a–c) Scores of the first two PCs, colored by life stage, with 95% confidence ellipses and intervals. (d) Cumulative total variance explained by the first ten PCs and variance of PC scores explained by life stage.

**Fig. S28.**
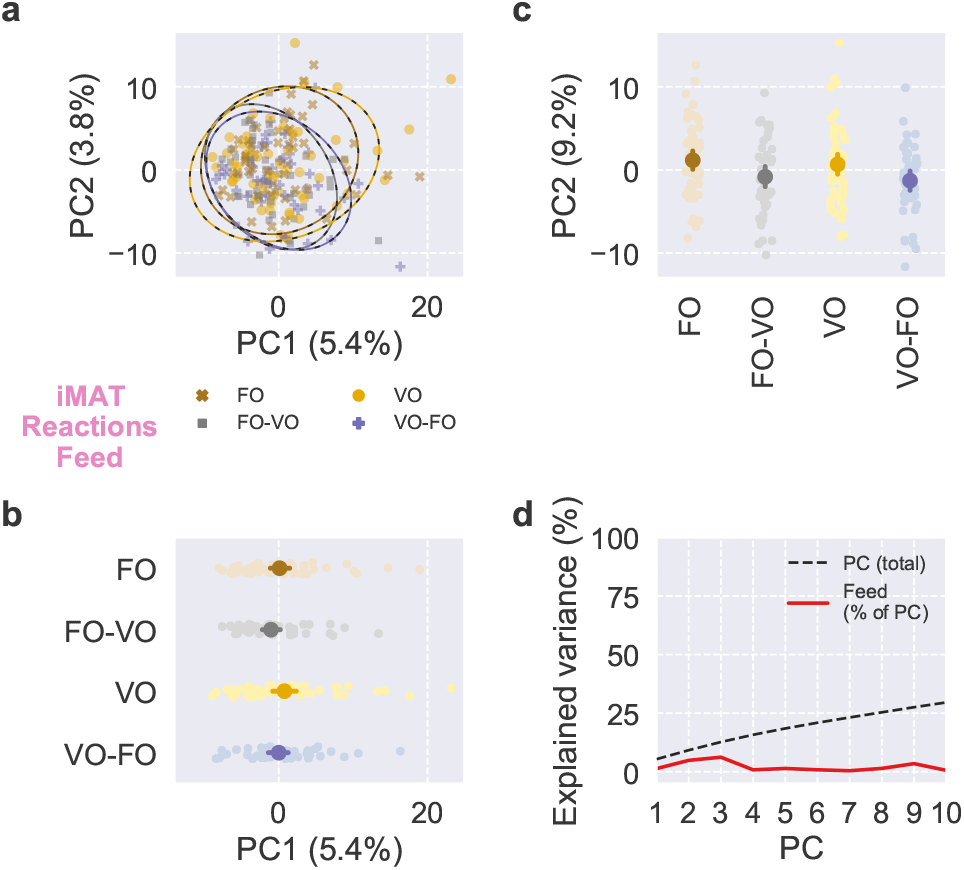
PCA of reaction presence for iMAT. Scores and explained variance from PCA of reaction presence for iMAT. (a–c) Scores of the first two PCs, colored by feed, with 95% confidence ellipses and intervals. (d) Cumulative total variance explained by the first ten PCs and variance of PC scores explained by feed.

**Fig. S29.**
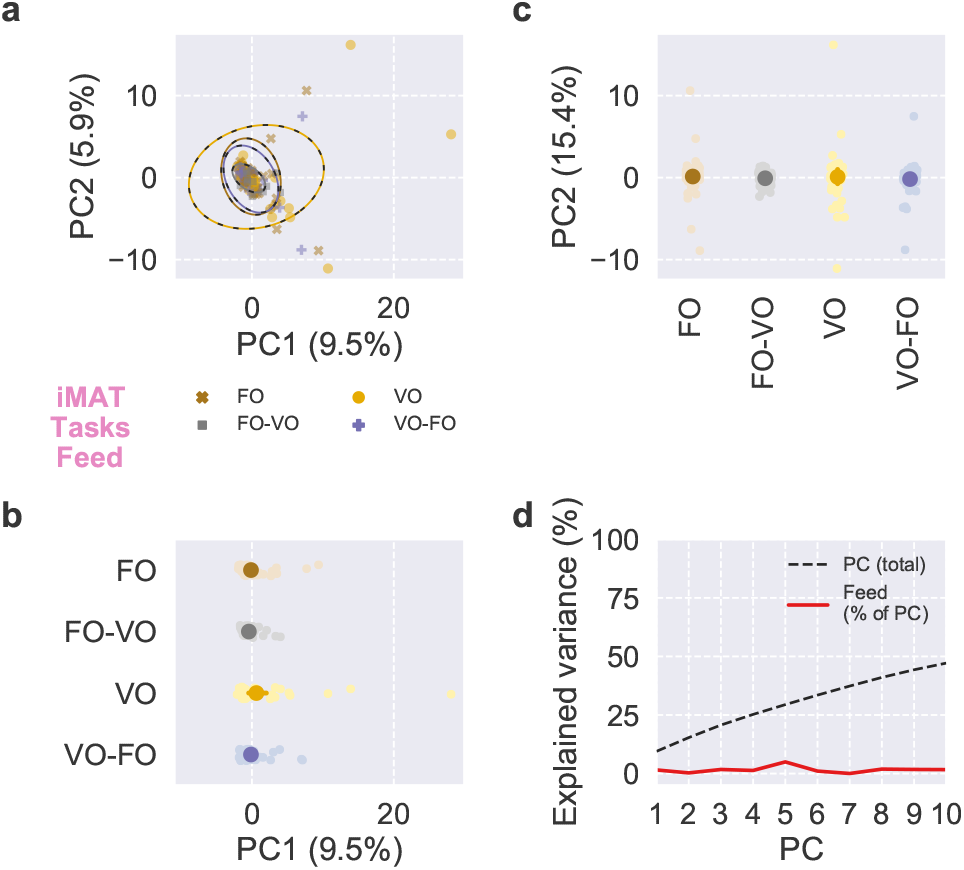
PCA of task feasibility for iMAT. Scores and explained variance from PCA of task feasibility for iMAT. (a–c) Scores of the first two PCs, colored by feed, with 95% confidence ellipses and intervals. (d) Cumulative total variance explained by the first ten PCs and variance of PC scores explained by feed.

**Fig. S30.**
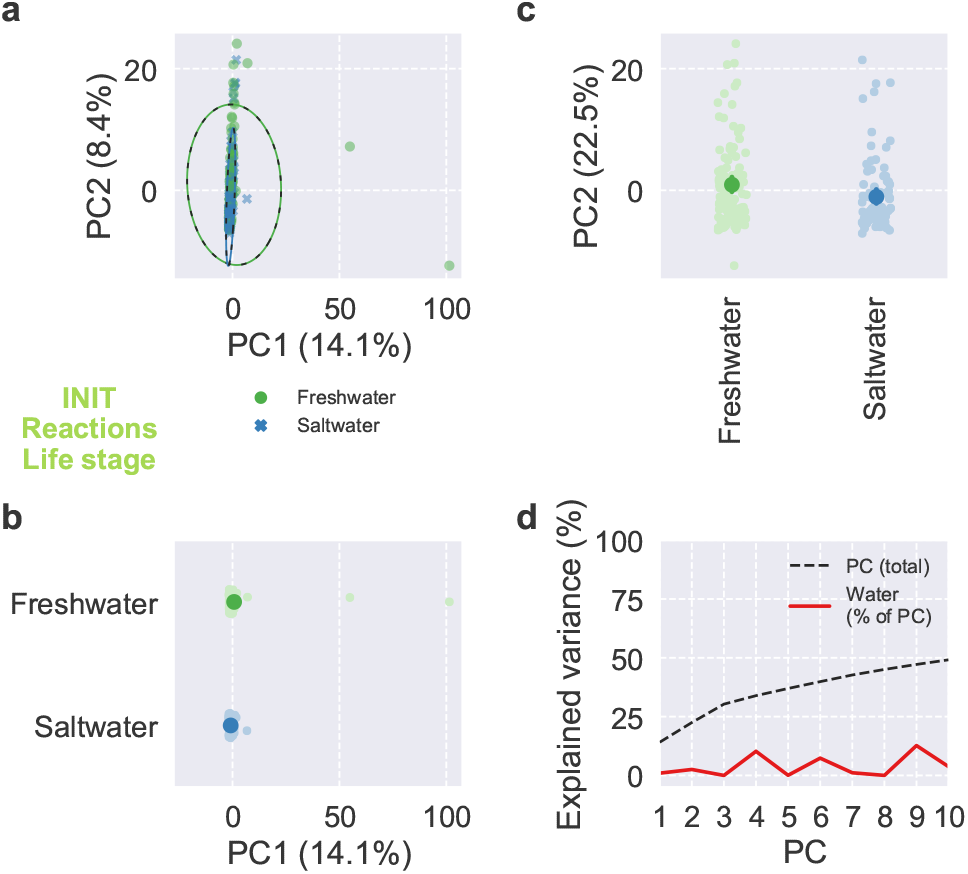
PCA of reaction presence for INIT. Scores and explained variance from PCA of reaction presence for INIT. (a–c) Scores of the first two PCs, colored by life stage, with 95% confidence ellipses and intervals. (d) Cumulative total variance explained by the first ten PCs and variance of PC scores explained by life stage.

**Fig. S31.**
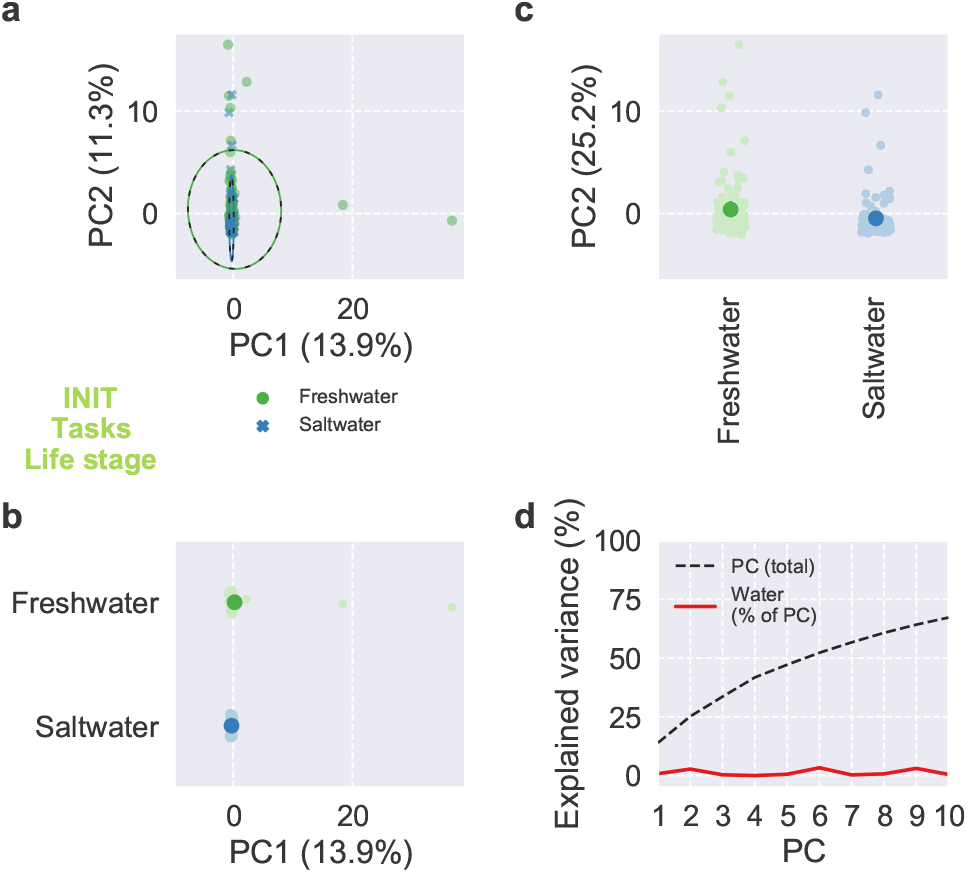
PCA of task feasibility for INIT. Scores and explained variance from PCA of task feasibility for INIT. (a–c) Scores of the first two PCs, colored by life stage, with 95% confidence ellipses and intervals. (d) Cumulative total variance explained by the first ten PCs and variance of PC scores explained by life stage.

**Fig. S32.**
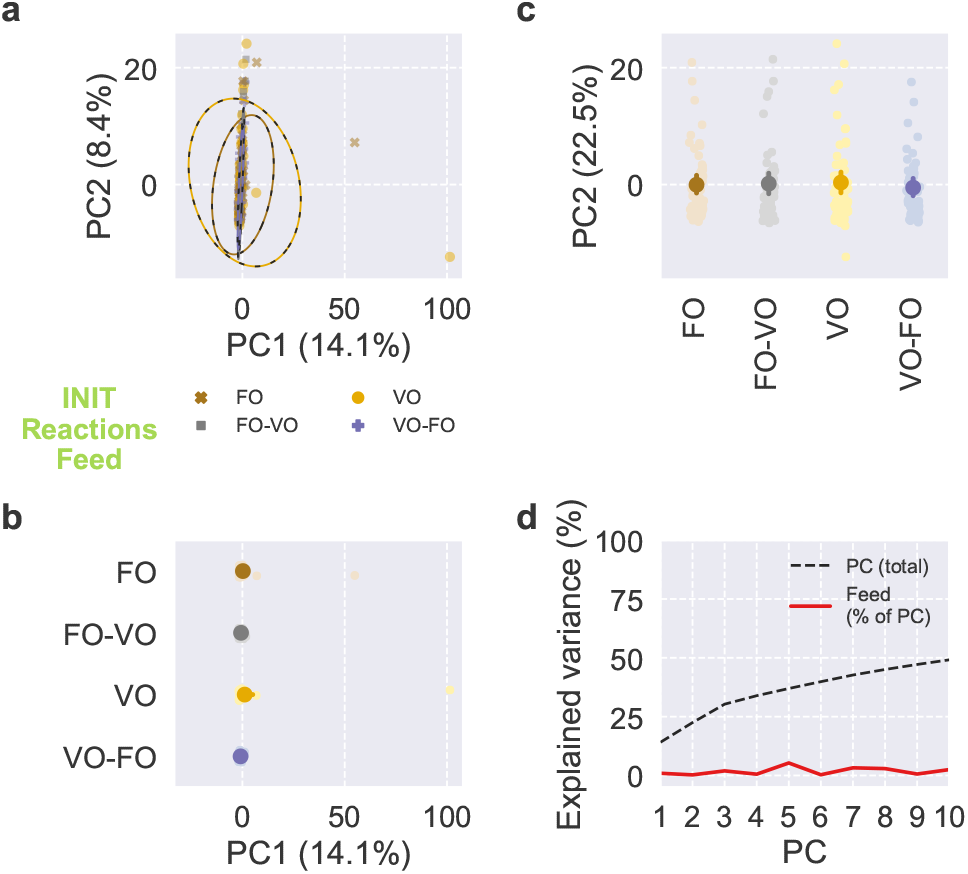
PCA of reaction presence for INIT. Scores and explained variance from PCA of reaction presence for INIT. (a–c) Scores of the first two PCs, colored by feed, with 95% confidence ellipses and intervals. (d) Cumulative total variance explained by the first ten PCs and variance of PC scores explained by feed.

**Fig. S33.**
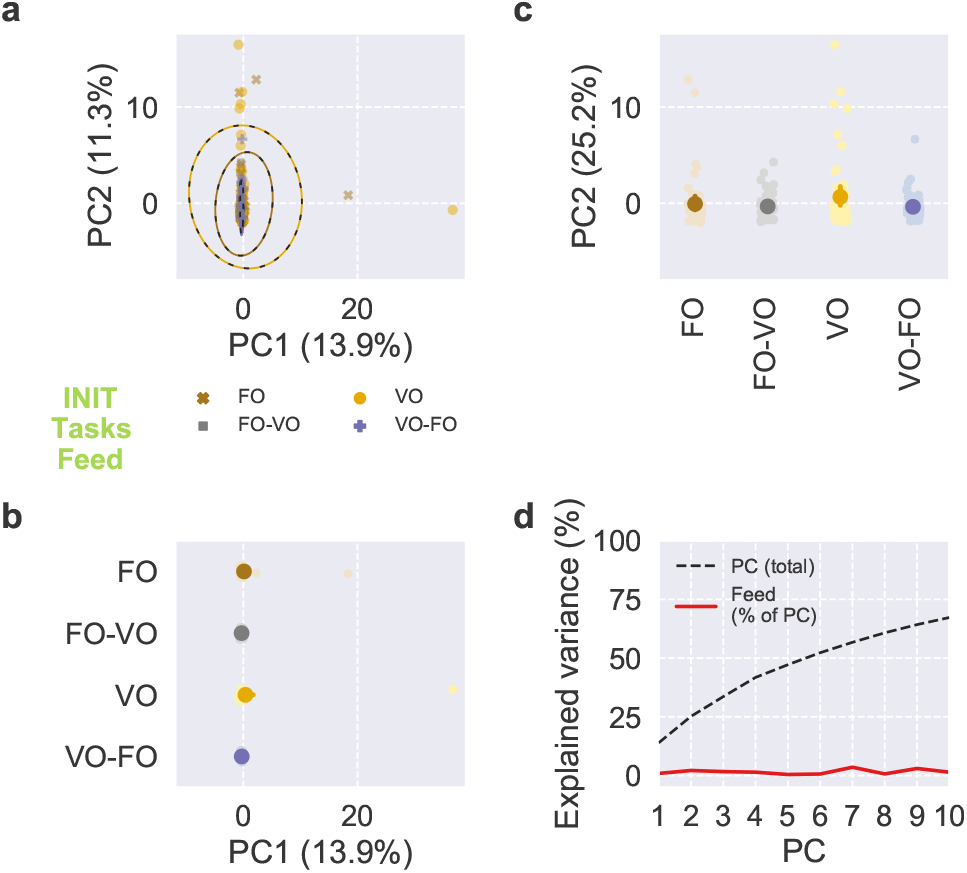
PCA of task feasibility for INIT. Scores and explained variance from PCA of task feasibility for INIT. (a–c) Scores of the first two PCs, colored by feed, with 95% confidence ellipses and intervals. (d) Cumulative total variance explained by the first ten PCs and variance of PC scores explained by feed.

**Fig. S34.**
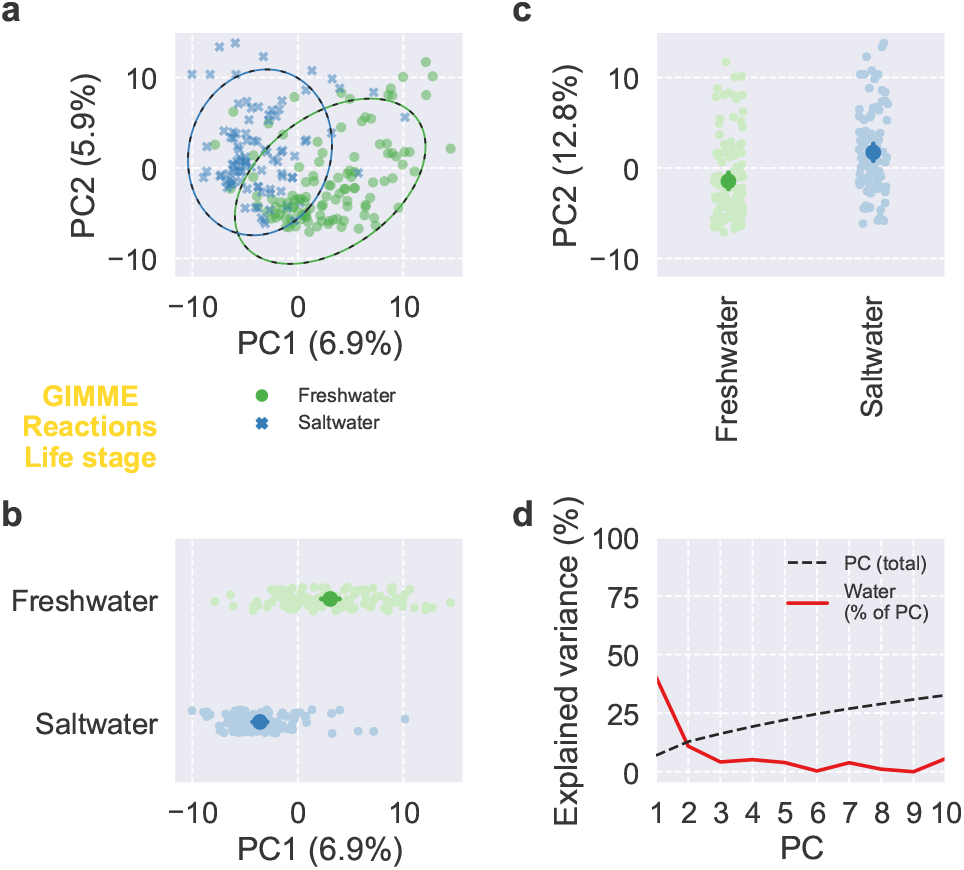
PCA of reaction presence for GIMME. Scores and explained variance from PCA of reaction presence for GIMME. (a–c) Scores of the first two PCs, colored by life stage, with 95% confidence ellipses and intervals. (d) Cumulative total variance explained by the first ten PCs and variance of PC scores explained by life stage.

**Fig. S35.**
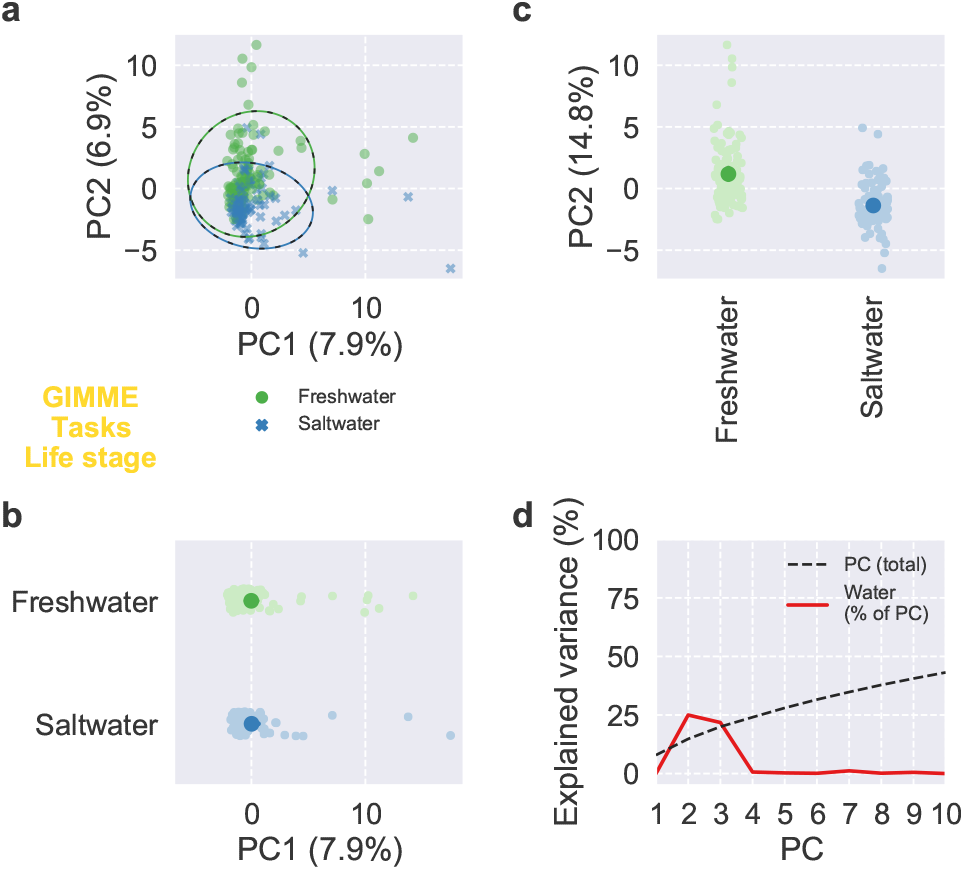
PCA of task feasibility for GIMME. Scores and explained variance from PCA of task feasibility for GIMME. (a–c) Scores of the first two PCs, colored by life stage, with 95% confidence ellipses and intervals. (d) Cumulative total variance explained by the first ten PCs and variance of PC scores explained by life stage.

**Fig. S36.**
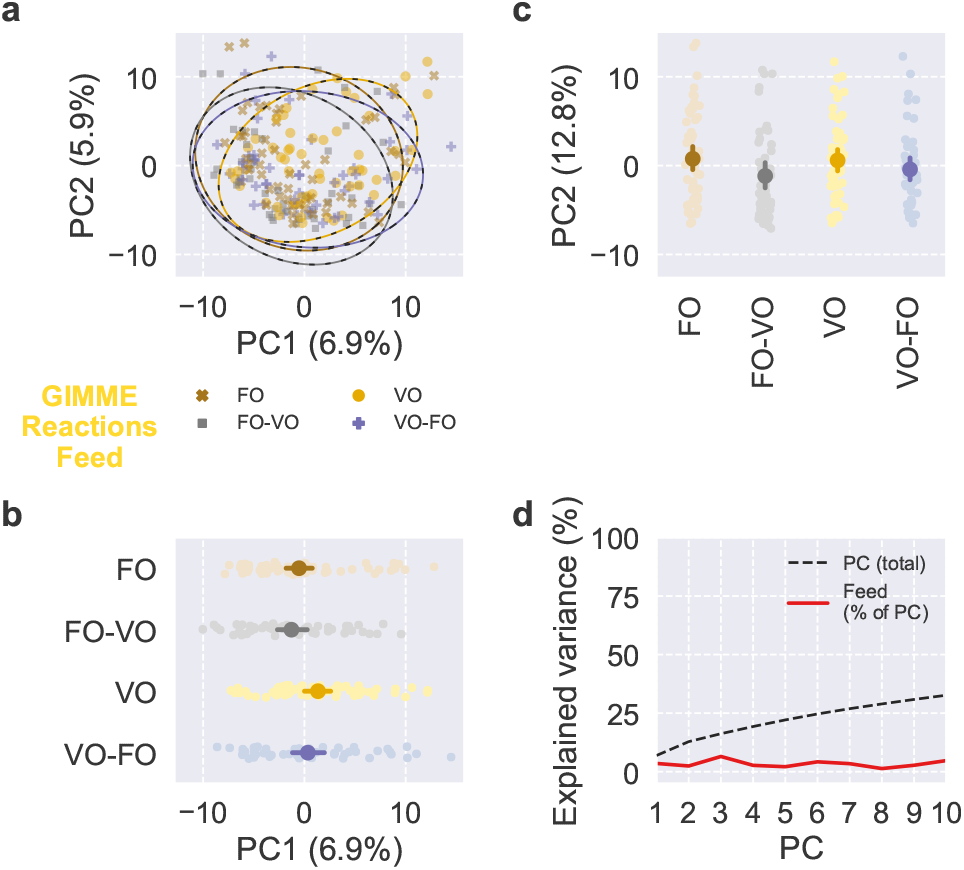
PCA of reaction presence for GIMME. Scores and explained variance from PCA of reaction presence for GIMME. (a–c) Scores of the first two PCs, colored by feed, with 95% confidence ellipses and intervals. (d) Cumulative total variance explained by the first ten PCs and variance of PC scores explained by feed.

**Fig. S37.**
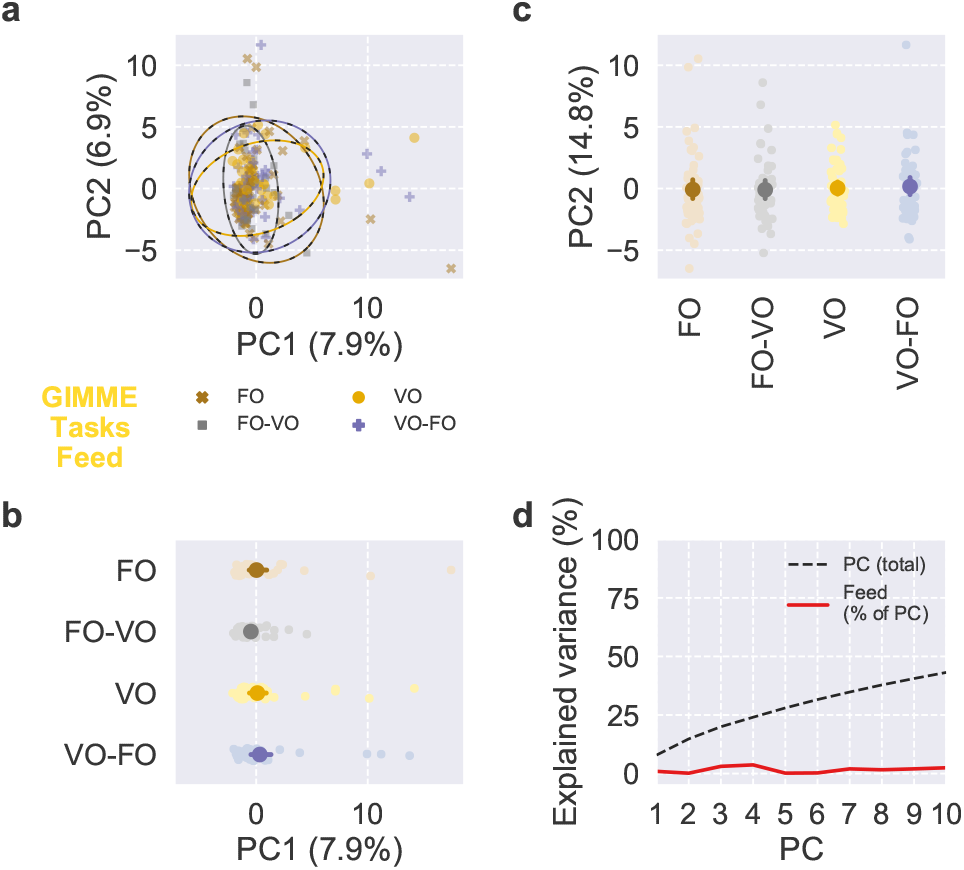
PCA of task feasibility for GIMME. Scores and explained variance from PCA of task feasibility for GIMME. (a–c) Scores of the first two PCs, colored by feed, with 95% confidence ellipses and intervals. (d) Cumulative total variance explained by the first ten PCs and variance of PC scores explained by feed.

**Fig. S38.**
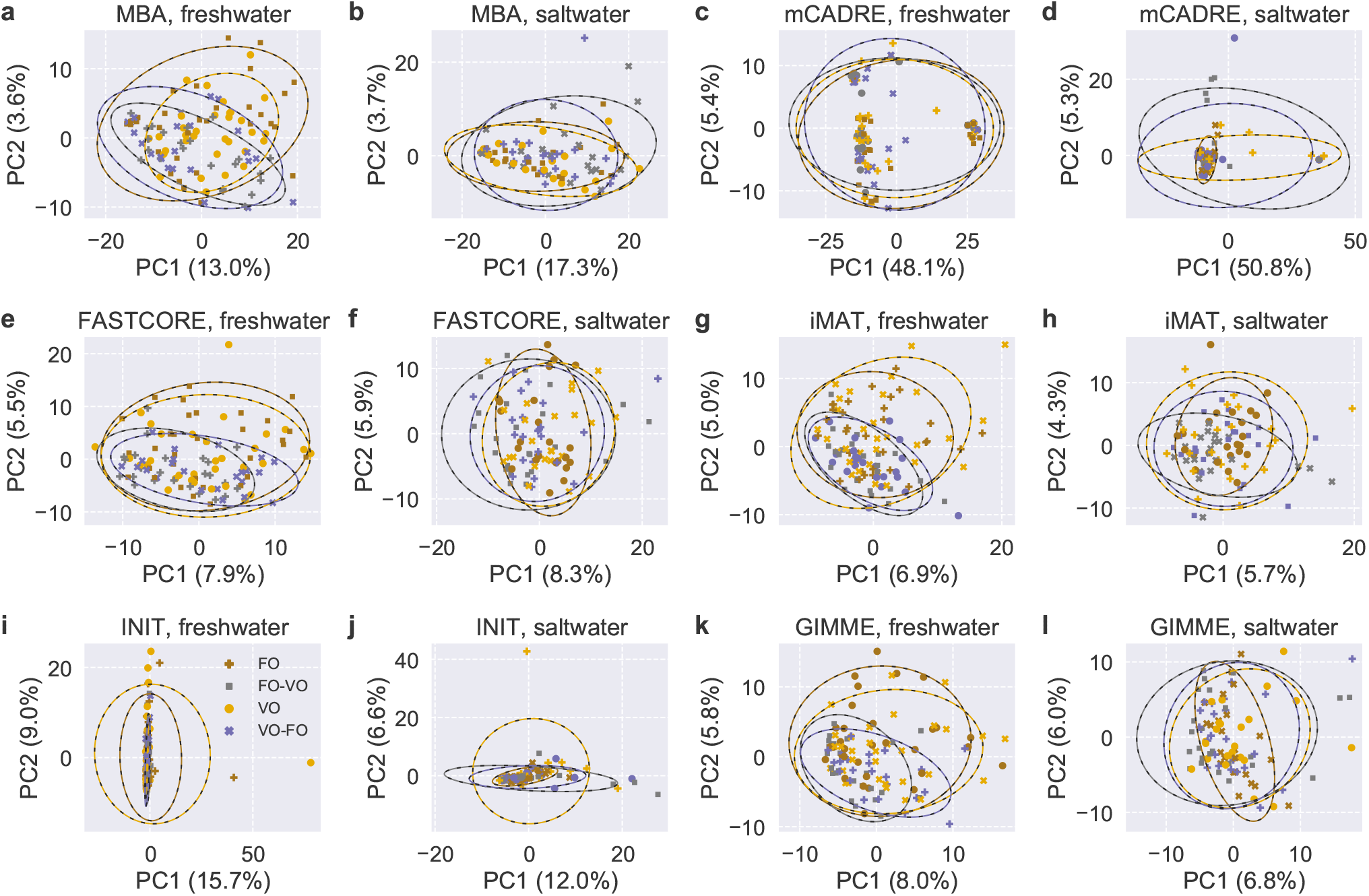
PCA of reaction presence and task feasibility within MEMs and life stages. Scores and explained variance of the first two PCs are shown, with colors indicating feeds and 95% confidence ellipses.

**Fig. S39.**
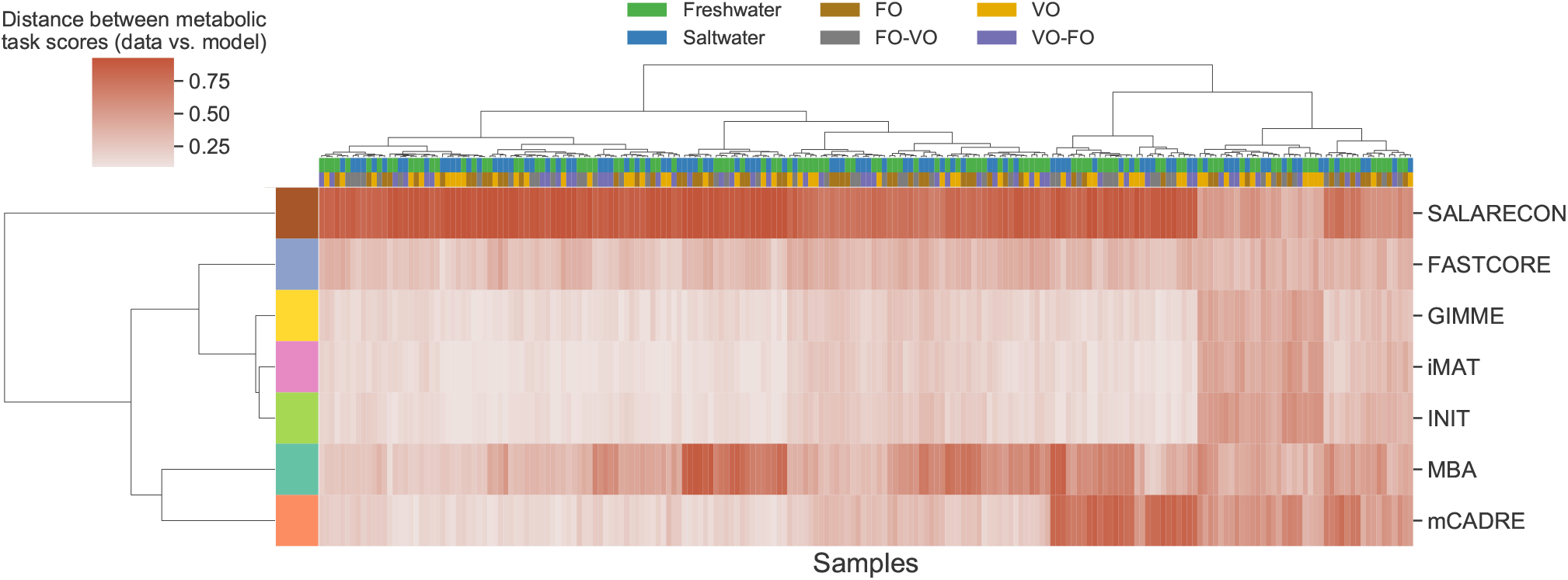
Clustered heatmap of metabolic task distances. Rows are MEMs, columns are models, and cells indicate the Normalized Hamming distance between task scores inferred from data and task scores predicted by models. Rows and columns are clustered by Ward’s minimum variance method, rows are colored by MEM, and columns are colored by life stage and diet.

